# An epigenetic basis for sustained inflammatory epithelial progenitor cell states in Crohn’s disease

**DOI:** 10.1101/2024.11.22.622113

**Authors:** Tatiana A. Karakasheva, Clara Morral Martinez, Yusen Zhou, Jingya Qui, Xinyi E. Chen, Gloria E. Soto, Shaneice K. Nettleford, Olivia T. Hix, Daana M. Roach, Alyssa M. Laguerta, Anusha Thadi, Rachael M. Edwards, Daniel Aleynick, Sarah Weinbrom, Elizaveta Borodyanskaya, Oliver H. Pickering, MaryKate Fulton, Chia-Hui Chen, Bella V. Peterson, Erik B. Hagen, Ian P. Yannuzzi, Zainab Haider, Zvi Cramer, Maire Conrad, Ning Li, Meenakshi Bewtra, Yasin Uzun, Kai Tan, Judith R. Kelsen, Andy J. Minn, Christopher J. Lengner, Kathryn E. Hamilton

## Abstract

Defining consequential differences in intestinal epithelial stem cells in healthy humans versus those with inflammatory bowel disease (Crohn’s disease and ulcerative colitis) is essential for the development of much needed therapies to restore the epithelial barrier and maintain its fidelity. Employing single cell transcriptomic/epigenomic approaches and colonoid models from children and adults with Crohn’s disease led us to identify an inflammatory secretory progenitor (ISP) cell state present almost exclusively in patients with Crohn’s disease compared to control subjects. ISPs exhibit gene expression profiles consistent with normal secretory progenitor cells but concomitantly express a suite of distinguishing pro-inflammatory genes. Mechanistically, ISPs exhibit open chromatin and gene expression of ISP signature genes. While these ISP-specific genes are not expressed in intestinal stem cells, their chromatin is accessible in Crohn’s disease stem cells suggesting that ISP genes are epigenetically poised in stem cells and are transcriptionally activated in ISP cells in the presence of inflammatory stimuli. Consistently, Crohn’s disease colonoids exhibit sustained ISP gene expression that can be elicited further with pro-inflammatory cytokines or via co-culture with pro-inflammatory macrophages. In summary, we define differences in the epithelial stem and progenitor compartment of patients with Crohn’s disease suggesting aberrant stem cell differentiation and inflammatory gene expression arises during disease.

**HIGHLIGHTS:**

- We identify an inflammatory secretory progenitor cell state in endoscopically non-inflamed tissue from pediatric and adult patients with Crohn’s disease.
- Crohn’s disease epithelial stem and progenitor cells display increased chromatin accessibility preceding the emergence of inflammatory secretory progenitor cell states.
- Crohn’s disease colonoids from non-inflamed regions contain inflammatory secretory progenitor cells in the absence of inflammatory stimuli
- Inflammatory secretory progenitor cells expand in colonoid culture in response to cytokine stimulation or co-culture with pro-inflammatory macrophages

## INTRODUCTION

The gastrointestinal tract comprises a diverse population of cells including epithelial, immune, and stromal cells, which together maintain intestinal barrier integrity. Epithelial cells are the arbiters between gut luminal contents and the underlying immune cells, promoting a dynamic barrier for organismal health. Failure of barrier restoration post-damage is a hallmark of inflammatory bowel disease (IBD, e.g. Crohn’s disease, ulcerative colitis, and IBD undefined)^1–3^, the incidence of which is increasing in adults, but even more so in children^4,5^. Children with IBD often experience more severe disease compared to adults, including an increased risk for surgery in pediatric patients^6^. Recent studies in patient tissue have advanced our understanding of cell type heterogeneity in patients with IBD^7–12^, however, epithelial stem cell function has not been thoroughly characterized in these patients. Prior studies provide evidence for both homeostatic and regenerative (i.e., facultative) epithelial stem cells in human tissues, but it remains unknown how chronic inflammatory environments functionally alter these stem cells.

Epithelial stem cells driven by canonical WNT pathway activity contribute to homeostatic maintenance of the epithelial barrier. In the context of tissue injury when WNT-driven, proliferative stem cells are damaged or killed, a subset of other epithelial cells exhibit plasticity that enables a pro-regenerative state to restore the epithelial barrier^11,13–18^. The reactivation of fetal intestinal gene expression programs is associated with regeneration from facultative stem cell populations; however, studies demonstrating these paradigms were performed in mouse models. Little is known about how epithelial stem cell populations are affected by chronic inflammation, such as in human IBD. Furthermore, mechanisms underlying changes in stem cells in human disease are not well defined.

It is well known that epigenetic modifications contribute to stem cell fate determination. For example, prior studies in mice show increased DNA methylation of stem cell genes and decreased methylation of mature lineage genes during the transition from stem to differentiated cells ^19,20^. However, recent evidence also highlights disease-driven alterations to the epigenome that are associated with chronic inflammation. Some of these epigenetic changes are thought to be a mechanism for inflammatory memory – an adaptive response tissues undergo after injury or inflammation. While inflammatory memory has been typically described in immune cells, recent studies in mice demonstrate that mammalian epithelial cells, particularly from progenitor compartments, can acquire enhanced transcriptional responses after an initial stimulus due to persistent open chromatin in tissue-specific memory domains^21–2711/20/2024^ 11:22:00 AM. While such inflammatory memory may be beneficial for enabling more rapid and robust immune responses to repeated pathogen exposure, the same phenomenon may contribute to the pathogenesis of chronic inflammatory disorders. Patient-derived organoid systems offer a unique opportunity to study inflammatory memory because they disentangle epithelial-intrinsic phenomena from those elicited by microenvironmental cues in the complex, inflamed in vivo environment. Work in human organoids suggests persistent epigenetic and functional differences in the IBD epithelium. For example, aberrant transcriptional signatures have been reported in tissue from IBD patients in remission^28,29^ and differential DNA methylation has been reported in organoids in patients with ulcerative colitis and Crohn’s disease^30^. Studies by our own group and others demonstrate a reduction in growth efficiency in colonoid cultures from patients with IBD that persists over numerous passages^31,32^. Finally, recent studies in mice with intestinal damage associated with graft-versus-host disease demonstrate that intestinal stem cell growth defects persist in culture, further supporting the notion that the intestinal stem cells can retain memory of past damage with negative consequences^33^. Whether common epigenetic mechanisms contribute to aberrant epithelial cell behavior in the context of chronic inflammatory diseases including Crohn’s disease is thus a question of significant interest.

In this study, we investigate how intestinal epithelial cells adapt to chronic inflammation that occurs in Crohn’s disease. Using endoscopic biopsies and colonoids from adult and pediatric subjects, we identify a disease-specific cell state, denoted the inflammatory secretory progenitor (ISP) state, possessing chromatin and transcriptional features consistent with inflammatory memory and pathologic gene expression. Epigenetic changes begin in stem and progenitor cells and are coupled to enhanced responsiveness to myeloid-secreted cytokines and impaired stem cell fitness. Taken together, our data support the notion that Crohn’s disease epithelial stem cells are epigenetically rewired to permit emergence of ISPs linked to features of disease pathology.

## RESULTS

### Emergence of inflammatory secretory progenitor cell states in Crohn’s disease epithelium

To map the differences in epithelial cell types and states between control and Crohn’s disease subjects, we collected biopsies from endoscopically non-inflamed areas of the ascending colon from 19 patients with Crohn’s disease (10 pediatric, 9 adult) and 23 control subjects (12 pediatric, 11 adult) (Supplemental Figure 1A; Supplemental Table 1). Using our optimized protocol to preserve live epithelial cells^34^, we acquired 106,583 unique transcriptomes (63,784 pediatric, 42,799 adult), which were separated into 12 clusters by unsupervised clustering analysis (Figure 1A, Supplemental Figure 1B,C): epithelium (38,045), CD4+ T cell (7,770), CD8+ T cell (12,644), B cell (18,248), plasma cell (15,135), macrophage (1,181), mast cell (982), endothelium (3,597), fibroblast (6,179), myofibroblast (745), pericyte (832), glia (1,225). The epithelium was further separated into 13 sub-clusters (Figure 1B, Supplemental Figure 1D-E, and Supplemental Table 2): stem cells (3,530), transit amplifying (TA) cells (2,170), early progenitors (9,004), *OLFM4+REG1A+* secretory progenitors (2,115), *OLFM4-REG1A+* secretory progenitors (947), *OLFM4-REG1A+* inflammatory secretory progenitors (905), absorptive colonocyte progenitors (5,701)), *FABP1+* absorptive colonocytes (2,500), *AQP8+* absorptive colonocytes (3,079), *BEST4+* colonocytes (830), goblet cells (1,010), tuft cells (286), and M cells (49). We observed differences in yield for epithelial sub-clusters in the pediatric Crohn’s disease cohort compared to pediatric controls, although only the decrease in BEST4+ cells and increase in *OLFM4+REG1A+* secretory progenitors reached statistical significance (P=0.01 and 0.05, respectively) (Supplemental Figure 1E). One cell cluster, which we designated *inflammatory secretory progenitors (ISP),* was predominantly present in Crohn’s disease and scarce in control samples (Figure 1C). ISPs lack markers of stem or TA cells, and express genes consistent with *OLFM4-REG1A+* secretory progenitors from healthy tissue, yet also express an inflammation-related signature. The top 10 ISP cluster markers are genes encoding antimicrobial peptides (*LCN2*, *REG1A*, *REG1B*, *DMBT1*, *PI3*), antigen presentation machinery (*CD74*, *HLA-DRA*, *HLA-DPA1*, *HLA-DMA*, *PSMB*), enzymes involved in mucosal immunity (*DUOX2*, *DUOXA2*, *ASS1*, *PLA2G2A*), and the chemokine *CCL20* (Figure 1D-F). Thus the disease-associated ISP cell state is characterized by enrichment for multiple genes previously linked to IBD^29,35–42,49^. We did not detect typical marker genes for Paneth or deep crypt secretory cells ^50,51^.

**Figure 1:**
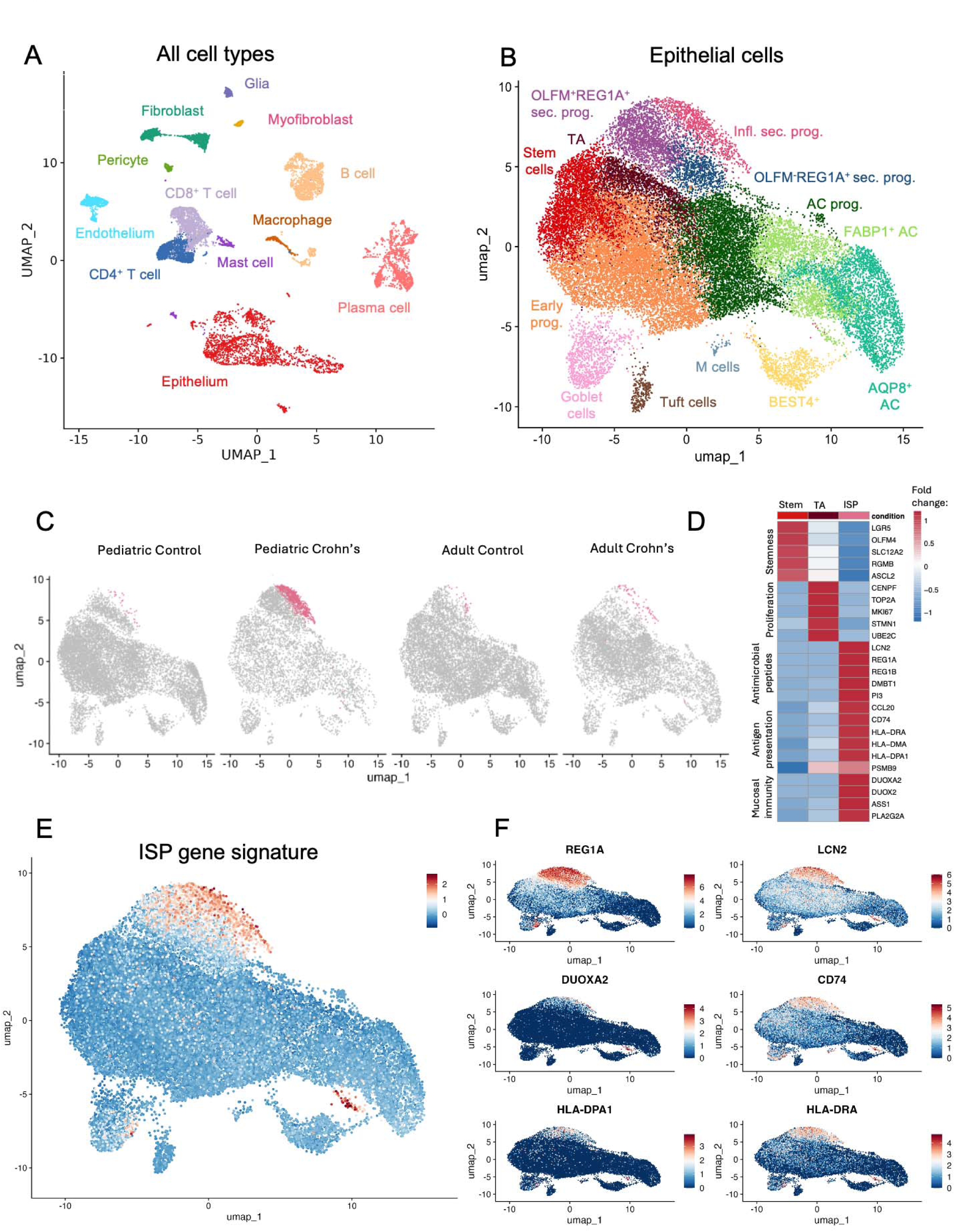
Inflammatory secretory progenitors constitute a cell state specific to Crohn’s disease. (A-B) UMAP visualization of all transcriptomes in whole biopsy (A) and sub-clustering of the epithelial transcriptomes (B). (C) UMAP projections of epithelial transcriptomes from pediatric control subjects, pediatric Crohn’s disease subjects, adult control subjects, adult Crohn’s disease subjects, with inflammatory secretory progenitor (ISP) cluster highlighted in color. (D) Heatmap summarizing relative expression of stem cell, TA, or ISP cluster marker genes (expression levels plotted averaged per cluster). (E-F) ISP gene signature enrichment and expression of individual ISP marker genes plotted on the epithelial UMAP. In all panels, N = 19 Crohn’s disease (10 pediatric, 9 adult) and 23 control subjects (12 pediatric, 11 adult).

### The inflammatory secretory progenitor cell state is associated with disease activity

Further evaluation of ISP abundance within the epithelial cluster demonstrated that there were significantly more ISPs in pediatric patients with active Crohn’s disease (pediatric Crohn’s disease activity index^52,53^, PCDAI>10) compared to patients with quiescent disease (PCDAI≤10) or control subjects (Figure 2A). This likely explains why ISPs were less prevalent in our adult cohort, since 88% of adult patients with Crohn’s disease were in remission at the time of colonoscopy (Supplemental Table 1; Supplemental Figure 2A). In addition to our cohorts, we observed ISPs in multiple publicly available datasets using reference-based integration, including pediatric and adult Crohn’s disease, as well as ulcerative colitis^8^, suggesting a common mechanism of ISP cell state induction across intestinal inflammatory diseases^3^. When comparing samples from inflammation-adjacent regions to samples from actively inflamed regions in these published datasets, ISPs were more abundant in inflamed tissue^12^ (Figure 2B, Supplemental Figure 2B-D). To validate the presence of ISPs in tissue, we stained ascending colon biopsies from the same patients as were sequenced (areas adjacent to inflammation) for hallmark ISP proteins LCN2, CD74, and HLA-DR. No co-staining of epithelium with all three markers was detected in control tissue, while in Crohn’s disease tissue we detected concurrent epithelial expression of all markers (Figure 2C, D, and Supplementary figure 2E). The abundance of HLA-DR^+^LCN2^+^CD74^+^ cells in tissue sections correlated with the abundance of ISP cells identified for the same individual subjects via scRNA-seq (Figure 2E).

**Figure 2:**
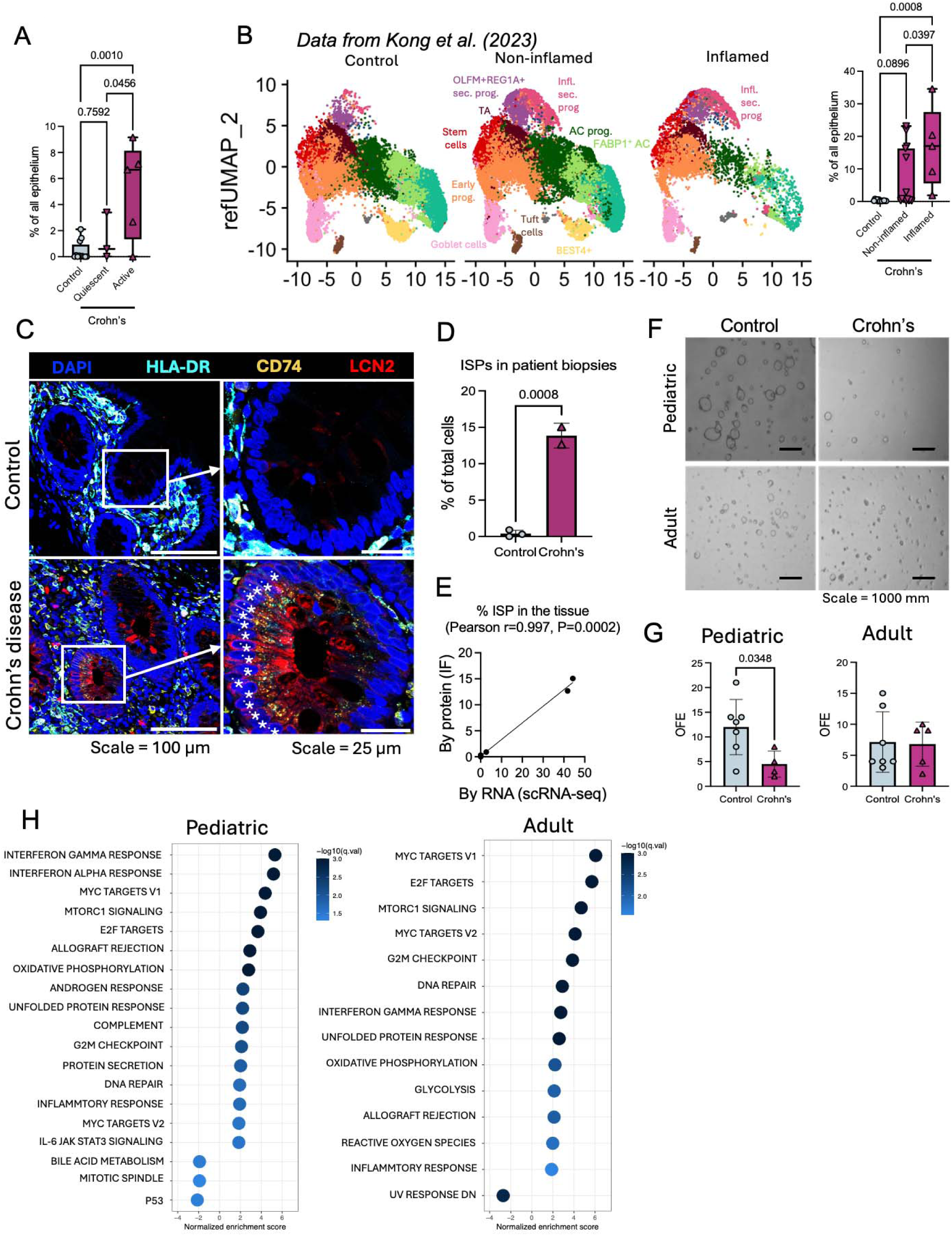
Inflammatory secretory progenitor state is a characteristic of active disease and is associated with loss of lineage fidelity. (A) Relative abundance of the ISP cluster in total epithelium for individual pediatric subjects (in active Crohn’s disease, biopsies were taken from areas adjacent to inflammation), P values (ordinary one-way ANOVA) presented on the plots. (B) UMAP projections on a published dataset (DUOS-000145, DUOS-000146; adult Crohn’s disease) and quantification of relative ISP abundance on the right P values (ordinary one-way ANOVA) presented on the plots. (C) Representative images of immunostaining for LCN2, CD74, HLA-DR in biopsies from study subjects. Asterisk denotes an ISP, quantified in (D) (N=2 subjects per group, X ROI analyzed per biopsy, and the average per subject plotted on graph). (E) Correlation analysis comparing the abundance of ISPs in tissue as quantified at transcript level (scRNA-seq) to quantification at protein level (immunostaining). (F) Representative images of colonoid cultures seeded as single cells to assess organoid formation efficacy (OFE). Scale, 100 um. OFE quantified in (G) as [N (colonoids formed)/N (cells seeded)]x100% (N= 7 pediatric controls, 4 pediatric Crohn’s; 7 adult controls, 5 adult Crohn’s; P=1-2). (H) HALLMARK pathways differentially enriched in stem cells from Crohn’s disease biopsies, compared to controls. Pathways with q<0.05 are presented.

We observed reduced stem cell abundance in pediatric Crohn’s disease (Supplemental Figure 1E), compelling us to evaluate how self-renewal in Crohn’s subjects is affected. Clonal organoid forming efficiency (OFE) assays demonstrated reduced OFE from pediatric Crohn’s disease subjects across multiple passages compared to control subjects, consistent with stem cell self-renewal deficits. We did not observe a difference in OFE between adult Crohn’s or control colonoids (Figure 2F-G). While reduced OFE was more prominent in pediatric compared to adult Crohn’s disease samples, there were significant gene expression changes in disease versus control stem cells in both cohorts: 87 genes in pediatric and 289 genes in adult stem cells were differentially expressed (Padj.<0.05). Furthermore, gene set enrichment analysis (GSEA) revealed upregulation of 16 and 13 pathways, mostly related to inflammation (and downregulation of 3 and 1 pathways) in stem cells from pediatric and adult Crohn’s, respectively (Figure 2H). Pathways related to cell cycle (Myc targets, E2F targets, G2M checkpoint) were most significantly upregulated in Crohn’s disease stem cells in our adult dataset. We considered the possibility that this finding could be a result of adult stem cells having higher activity of the above pathways, compared to pediatric stem cells. Upon comparison of differentially expressed genes, “Myc targets” pathway was the only cell cycle-related pathway that was upregulated in pediatric stem cells, compared to adult stem cells. These data together suggest that in subjects with active Crohn’s disease, stem cells from non-inflamed areas exhibit reduced stem cell function.

Recent studies in mice demonstrate the reactivation of a fetal intestinal gene expression program during post-injury repair in the adult. This program, sometimes referred to as a revival stem cell signature^54^ or fetal reprogramming signature^16,55^, is driven primarily by YAP activity and also IFNgamma^15,56^. In addition, pediatric Crohn’s disease epithelial cells share transcriptional profiles with human fetal intestinal epithelium^11^. Interestingly, “Interferon gamma response” was among the most significantly enriched gene sets in Crohn’s disease stem cells compared to control subjects (Figure 2H). We therefore interrogated our datasets for evidence of fetal gene expression programs. We found that ISPs were enriched for all signatures tested, while there was no difference between Crohn’s disease and control subjects in other cell clusters (Supplemental Figure 3A). These findings demonstrate that regenerative gene signatures persist in cells in the ISP state from non-inflamed Crohn’s disease tissue.

### Epigenetic memory underlies persistent inflammatory gene expression in Crohn’s disease epithelium

To evaluate how Crohn’s disease alters the epigenetic landscape of epithelial cells, including ISPs, we performed single-nucleus assay for transposase-accessible chromatin sequencing (snATAC-seq) on a subset of the same samples analyzed above. We analyzed chromatin accessibility of the epithelium from pediatric subjects since our pediatric Crohn’s cohort had higher representation of ISPs. Unsupervised cell clustering based on genome-wide chromatin accessibility patterns identified 12 distinct epithelial clusters from 5,541 epithelial cells distributed across 4 control (3,628) and 6 Crohn’s (1,913) pediatric samples after quality control filtering (Supplemental Figure 3B). To annotate these 12 clusters, we examined the inferred gene activity of published intestinal epithelial gene signatures^57^. We also included a custom ISP gene signature from our scRNA-seq analysis (Supplemental Table 4). Among all 12 clusters, clusters 2 and 3 showed specific enrichment for open chromatin at ISP signature genes but not for any other canonical intestinal gene programs; thus, we labeled these clusters as ISPs (Supplemental Figure 3C-E). Importantly, we also observed modest chromatin accessibility and gene activity enrichment of the ISP signature in the undifferentiated stem and progenitor cell clusters (Supplemental Figure 3E). Next, we examined changes in cellular composition between controls and Crohn’s disease samples. Consistent with what we found in the scRNA-seq dataset, ISP cells were specifically enriched in Crohn’s samples (P=0.098) at the expense of absorptive colonocytes (P=0.019) (Figure 3A, Supplemental Figure 3F).

**Figure 3:**
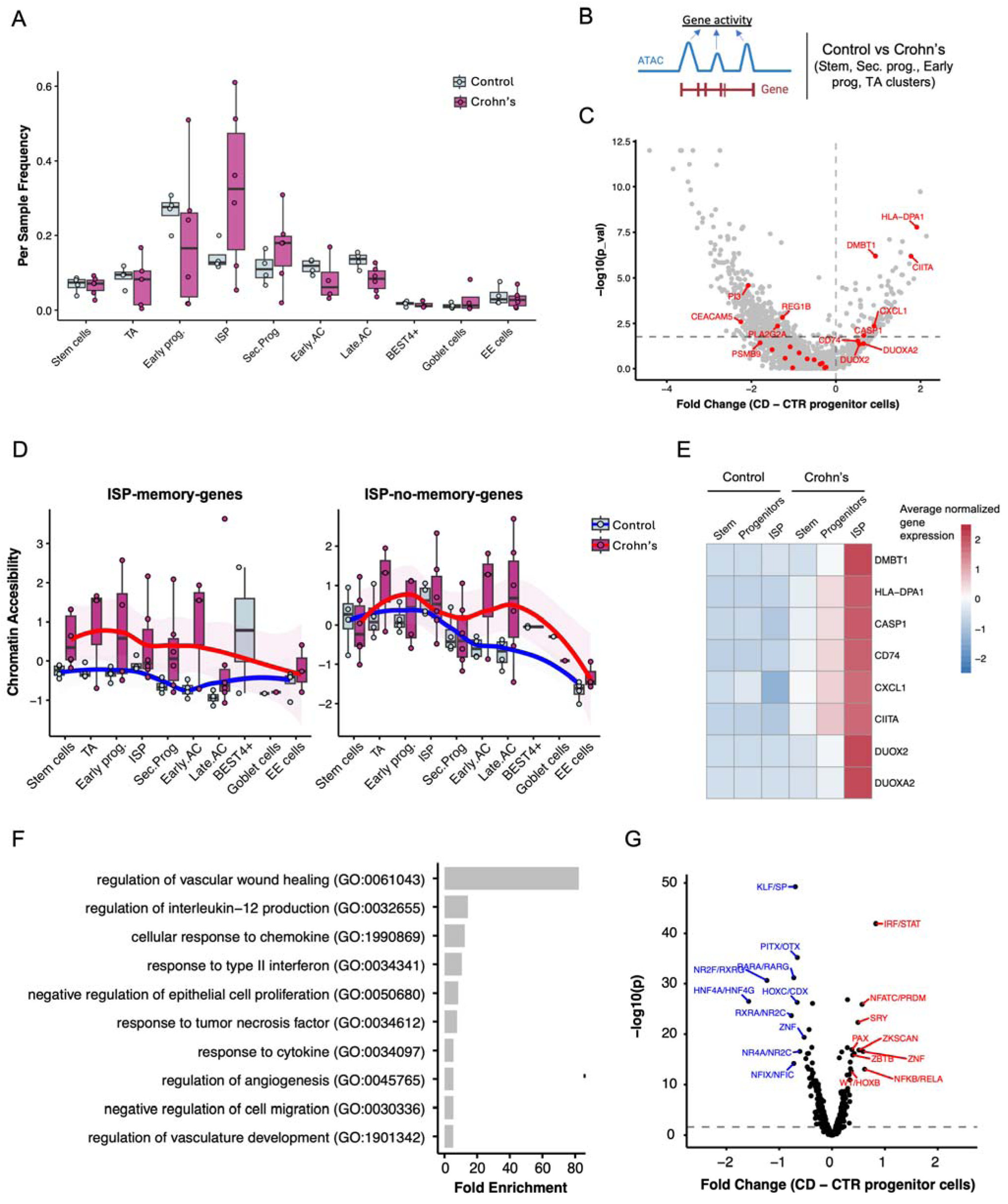
Chromatin accessibility is enhanced in a subset of ISP genes in Crohn’s disease epithelium. (A) Frequency of cell types comparing controls and Crohn’s patients. The boxplot represents the data’s interquartile range (IQR), with the median indicated. The whiskers represent the highest and lowest values within 1.5 times the IQR. The points represent individual sample frequencies, P values ≤ 0.1 presented on the plot. (B) Schematic representation of Gene activity score. (C) Volcano plot showing the differentially expressed genes in a merged Progenitor cluster (Stem, Secretory progenitors, Early progenitors, TA) between Crohn’s and control patients. The genes that belong to the ISP gene signature are highlighted in red. (D) Average chromatin accessibility for the “ISP-memory" genes (*HLA-DPA1, CIITA, DMBT1, CD74, CXCL1, CASP1, DUOX2, DUOXA2*) and “ISP-no-memory" genes (*REG1A, REG1B, LCN2, PLA2G2A, PI3, SPINK1, IFITM3, MUC1, ASS1, CCL20, PSMB9, HLA-DMA, IFI27, CEACAM5, MUC4*) in epithelial clusters in Crohn’s or control samples. (E) Heatmap summarizing average normalized expression of “ISP-chromatin-accessible" genes in Stem, Progenitor, and ISP, clusters in control or Crohn’s disease. (F) GO term analysis showing signaling pathways enriched in Crohn’s merged Progenitor cluster compared to controls. (H) Volcano plots representing enriched motifs in the merged Progenitor cluster in Crohn’s compared to controls.

We next delved deeper into chromatin accessibility patterns associated with emergence of the ISP cell state in Crohn’s disease. We predicted differential gene activity using the accessibility profiles of chromatin regions near gene loci in cell types presumed to give rise to ISPs -stem cells, secretory progenitors, early progenitors, and TA cells - between Crohn’s versus control (Figure 3B). We found a total of 393 genes (out of 1858 highly variable genes) that are differentially accessible between conditions (Supplemental Table 5), including 8 of 23 hallmark ISP signature genes (D*MBT1*, *HLA-DPA1, CIITA, CD74, CXCL1, CASP1, DUOX2, and DUOXA2* (Figure 3C). Interestingly, some of these genes were associated previously with enhanced chromatin accessibility in other chronic inflammatory diseases^23,58,59^, suggesting that they represent common targets of epigenetic modification in response to chronic inflammation. These specific ISP genes that have higher chromatin accessibility in Crohn’s disease (*HLA-DPA1, CIITA, DMBT1, CD74, CXCL1, CASP1, DUOXA2, DUOX2*, which we term “ISP-memory” genes) may be a result of epigenetic memory that occurred *in vivo.* To investigate this hypothesis, we calculated average accessibility for these genes in epithelial sub-clusters compared to the remaining genes within the ISP signature (*REG1A, REG1B, LCN2, PLA2G2A, PI3, SPINK1, IFITM3, MUC1, ASS1, CCL20, PSMB9, HLA-DMA, IFI27, CEACAM5, MUC4*, or “ISP-no-memory” genes). The genes in the first group have enhanced accessibility in Crohn’s disease in stem cells (p = 0.0038) and persist through differentiated lineages, whereas in control samples these genes do not have enhanced chromatin accessibility. In contrast, the remaining ISP signature genes do not show enhanced chromatin accessibility in any progenitor clusters (Figure 3D). “ISP-memory” genes are expressed at lower levels in stem cells than in subsequent progenitor clusters and peak in the ISP cluster in Crohn’s disease but not control epithelium (Figure 3E), indicating that poised open chromatin precedes expression of these genes in Crohn’s disease.

In addition to ISP genes, there are 84 genes with enhanced accessibility in Crohn’s disease progenitor cell population compared to control. Gene ontology enrichment for these genes revealed several terms related to TNFalpha (“regulation of vascular wound healing”, “response to tumor necrosis factor”), as well as STAT (“response to type II interferon”, “response to cytokine”, “regulation of angiogenesis”, “regulation of vasculature development”) signaling, among the top enriched signaling pathways in Crohn’s patients compared to controls (Figure 3F). This is consistent with TNFalpha being a predominant cytokine found in Crohn’s patients^60^. The TNF cytokine family induces transcription of genes involved in several biological processes, such as inflammation, cell survival, and differentiation, primarily through activating the NFkB transcription factor. Accordingly, differential transcriptional factor binding analysis identified NFkB/RELA motifs as enriched in open chromatin peaks in Crohn’s disease samples (Figure 3G). These NFkB motifs showed specific enrichment in the stem cells and ISP cell clusters (Supplemental Figure 3G).

Since the motifs for NFkB/RELA, as well as interferon response factors (IRF)/STAT were enriched in the open chromatin of Crohn’s epithelia, we focused on ISP signature genes and investigated the distribution of chromatin accessibility peaks surrounding these regulatory regions. Looking at the level of progenitor cells (merged stem cells, TA, early progenitors, secretory progenitors, and ISP) or individual sub-clusters, we detected peaks associated with more open chromatin in Crohn’s disease for *HLA-DPA1*, *HLA-DRA*, *CD74*, and *DUOXA/DUOX2* genes (Figure 4A, Supplemental Figure 4), but not for *LCN2* gene (Supplemental figure 4). Interestingly, we observed subject-based variability in chromatin accessibility at the level of peak coverage across all peaks (Figure 4B). These data suggest that increased chromatin accessibility around pro-inflammatory transcription factor binding motifs may contribute to expression of ISP signature genes in Crohn’s disease epithelium and enable robust reactivation of these with reintroduction of inflammation. This, along with the increase in epigenetically-defined ISP clusters at the apparent expense of TA and early progenitor clusters, suggests a possible developmental relationship between ISPs and upstream cell populations.

**Figure 4:**
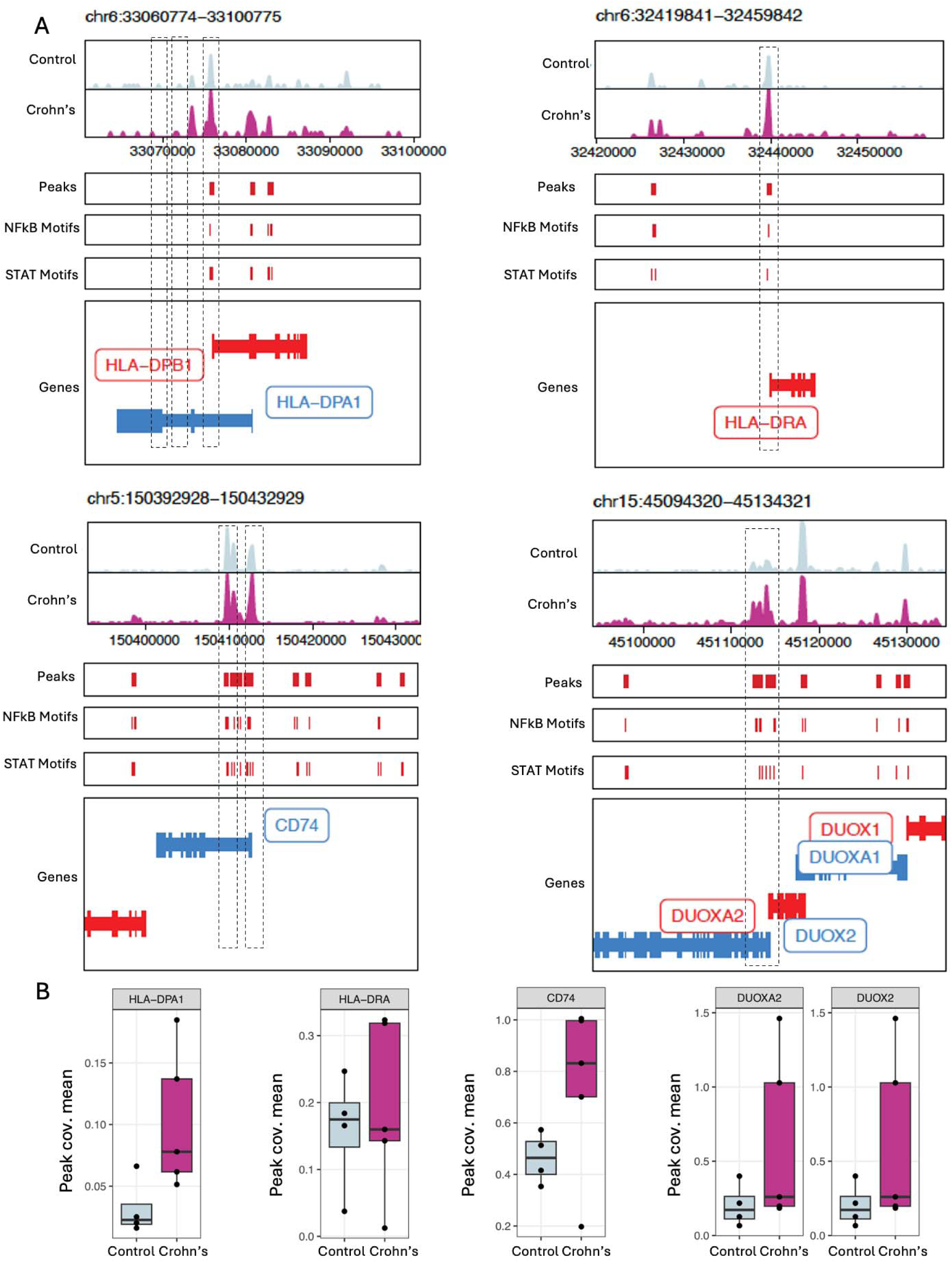
Select ISP signature genes contain open chromatin peaks enriched in Crohn’s disease. (A) Open chromatin peaks in the progenitor cells (merged stem cell, TA, early progenitors, secretory progenitors, ISP, clusters), with traces from control and Crohn’s disease subjects separated by color. Boxes placed to highlight the peaks with differential enrichment in Crohn’s vs control samples. Corresponding gene maps positioned below the chromatin traces. (B) Chromatin availability plotted as mean peak conversion values across all peaks, at subject level.

### Crohn’s disease colonoids are poised for induction of an ISP cell state

To test whether increased chromatin accessibility described above allows for Crohn’s epithelium to retain the ISP cell state, we turned to our colonoid bank derived from the same cell suspensions as were sequenced. Interestingly, we found that mRNA levels of ISP genes *HLA-DPA*, *HLA-DRA*, *CD74*, and *LCN2* were elevated in colonoids derived from Crohn’s disease subjects, compared to controls (Figure 5A). Yet, we did not detect equivalent staining for the corresponding proteins in colonoids at baseline (Figure 5B, Supplemental Figure 5A). Interestingly, we noticed that while there was no evidence of elevated HLA-DR expression on the surface of colonoids cultured in stem cell-favoring medium, when we induced differentiation, we found a significant (Pearson r=0.87, P=0.003) correlation between HLA-DR protein on the cell surface in colonoids and *HLA-DRA* mRNA counts in epithelial cell cluster by scRNA-seq of the biopsy from the corresponding study subject (Supplemental figure 5B). Thus, all experiments involving colonoids have been conducted in differentiation medium. We previously reported the need for cytokine re-stimulation for re-induction of HLA-DR protein expression in colonoids from very early onset IBD patients, despite the presence of detectable mRNA at baseline^32^. Therefore, we hypothesized that re-introduction of inflammatory stimuli may result in emergence of ISPs in vitro.

**Figure 5:**
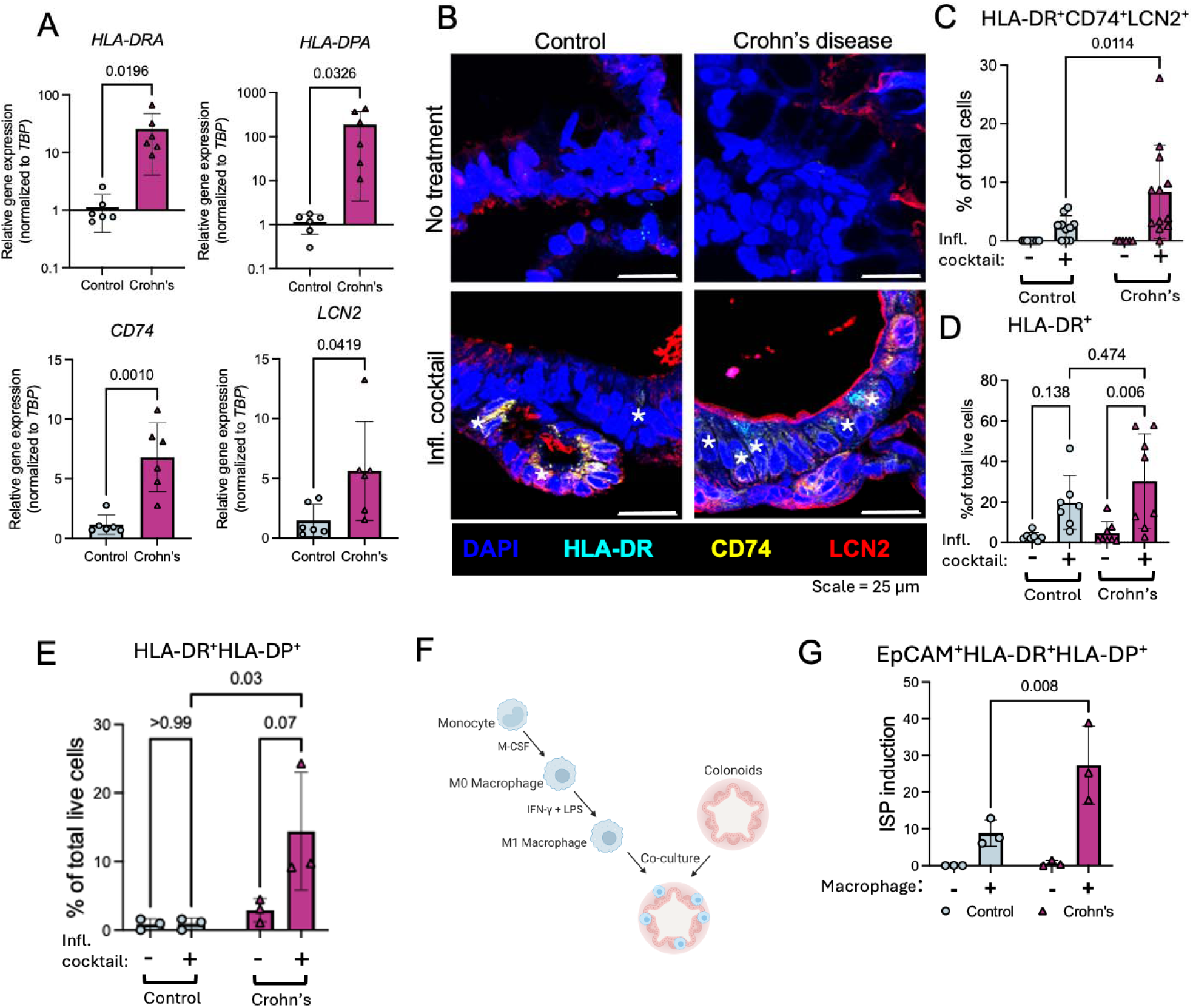
ISP state can be experimentally induced in patient-derived colonoids. (A) Expression of ISP marker genes HLA-DPA1, HLA-DRA, CD74, and LCN2, in colonoids from Crohn’s disease or control subjects (N=3 each, P=4-6). (B) Representative images of immunostaining for LCN2, CD74, HLA-DR in biopsies in colonoids from study subjects. Asterisk denotes an ISP. Quantified in (C): 2 lines (12 high power fields) per group, P=7-8. (D) Quantification (via flow cytometry) of % HLA-DR+ cells in colonoids treated with the inflammatory cocktail (20 ng/mL IL-1beta, 100 ng/mL TNFalpha, 1 ug/mL flagellin, and 10 ng/mL IL-6) for 24 hours (N= 7 or 8 lines from control subjects or Crohn’s disease patients, respectively, P=6-8). (E) Quantification (via flow cytometry) of % HLA-DR+HLA-DP+ cells in colonoids treated with the inflammatory cocktail for 24h. One line each from control subject or Crohn’s disease patient, N=3 independent experiments repeated in different passages (P=4-6). (F) Schematic: Monocytes from peripheral blood of healthy donors (different donor for each of the 3 independent experiments) were differentiated into M1 macrophages and embedded with colonoids (P4-7) from Crohn’s disease or control subjects (N=3 each), followed by flow cytometry 24 hours later. (G) Quantification of ISP induction (calculated as % EpCAM+HLA-DR+HLA-DP+ cells normalized to the degree of macrophage activation and polarization) in colonoid mono-cultures or co-cultures with M1 macrophages.

We employed a cocktail of 20 ng/mL IL-1beta, 100 ng/mL TNFalpha, 1 ug/mL flagellin, and 10 ng/mL IL-6, previously shown to induce IBD-relevant gene expression in colonoids^61,62^. Treatment of colonoids with this cytokine cocktail resulted in clusters of cells co-expressing ISP markers HLA-DR, LCN2, and CD74 in Crohn’s but not control colonoids (Figure 5B-C, Supplemental Figure 5A). By flow cytometry, inflammatory cocktail treatment resulted in potent upregulation of HLA-DR expression on the surface of cells in both control and Crohn’s disease colonoids, albeit the increase only reached statistical significance in Crohn’s disease colonoids (P=0.006, vs P=0.138 in control lines) (Figure 5D). We hypothesized that adding a second cell surface marker would detect a cell population selectively upregulated in response to the cocktail in Crohn’s colonoids only. Since our snATAC-seq data revealed that chromatin accessibility around the *HLA-DPA* gene was most potently increased in Crohn’s disease epithelium (Figure 3C), we used HLA-DP as a second marker to more specifically distinguish ISPs via flow cytometry. Indeed, cells co-expressing HLA-DP and HLA-DR were expanded significantly in colonoids from Crohn’s disease and not control subjects upon stimulation with the inflammatory cocktail (Figure 5E). To confirm that the HLA-DR^+^HLA-DP^+^ cell population is enriched for ISPs, we quantified expression of ISP marker genes *HLA-DPA*, *HLA-DRA*, *CD74,* and *LCN2,* in total live cells vs HLA-DR^+^HLA-DP^+^ cells sorted from colonoids – either untreated or subjected to the inflammatory cocktail for 24 hours (Supplemental Figure 5C). HLA-DR^+^HLA-DP^+^ cells were absent in control colonoids at baseline, and inflammatory cocktail treatment did not elicit an expansion sufficient to sort and extract RNA (Supplemental figure 5D). However, we found that HLA-DR+HLA-DP+ cells in untreated Crohn’s disease colonoids exhibit enrichment of hallmark ISP genes relative to total live cells (Supplemental figure 5E).

In search of a more physiological model of ISP induction, we turned to the immune microenvironment in colonic submucosa. Tissue-resident macrophages, especially M1-polarized, are known to secrete IL-1beta, TNFalpha, and IL-6, which we confirmed via cytokine bead array assay^63^ (Supplemental figure 5F). To test whether M1-polarized macrophages can induce an ISP cell state in vitro, we obtained human monocytes from healthy donors and differentiated them into macrophages ex vivo, followed by direct co-culture with control or Crohn’s disease colonoids (Figure 5F). We found that degree of macrophage differentiation varied based on the donor, with 41.5% to 85.2% of total CD14+ monocytes co-expressing macrophage markers CD64 and CD80 (Supplemental figure 5G-H) and normalized %ISP induction to the degree of macrophage activation in respective donor (see Methods). Interestingly, on average, colonoids from Crohn’sdisease subjects were 3x more responsive to ISP induction by M1 macrophages than control colonoids (Figure 5G). These findings suggest that in the mucosa of Crohn’s disease patients, the emergence of ISP cell state is driven at least in part by pro-inflammatory M1-polarized macrophages.

## DISCUSSION

Our study defines a new paradigm in which epigenetic memory promotes disease-associated stem cell changes that precede the emergence of cells in an ISP state in patients with IBD. While ISP marker genes have been broadly associated with IBD, their co-expression in a distinct progenitor population supports a new model in which pathogenic cell states may arise and persist in disease. Even more striking is the identification of ISPs in biopsies derived from uninflamed tissue. This suggests that the ISP state is either maintained by exposure to systemic inflammatory molecules or that the ISP state was acquired during active inflammation and that memory in the epithelial epigenome enables this state to persist in the absence of active inflammation at the tissue site. Analyses of increased chromatin accessibility around proinflammatory transcription factor binding sites of ISP marker genes support a mechanism by which these cells are poised to respond more robustly to subsequent inflammation. Data from human colonoids functionally validate that ISP marker gene expression remains elevated in homeostatic culture and that cells in this state are robustly expanded following inflammatory stimuli in Crohn’s epithelium. The fact that elevated cell expression of HLA-DA in unstimulated colonoids is only evident upon differentiation implies a role for the differentiated cells in sustaining the ISP state. Finally, data demonstrating reduced organoid formation from subjects whose epithelium harbors cells in the ISP state suggest that having cells in an ISP state may be detrimental to the reestablishment of the epithelial barrier post-injury. Taken together with differential chromatin accessibility findings, our data support the notion that an altered epigenome persists in non-inflamed areas that are primed to respond to subsequent inflammatory insults by producing cells in a pathogenic inflammatory secretory progenitor cell state.

Our finding that class II antigen presentation proteins HLA-DR and HLA-DP are specific markers for ISPs, including in vitro, lend support to the hypothesis that persistent ISP states are detrimental since deletion of MHCII in mouse models of colitis is associated with reduced mucosal inflammation^64^. Conversely, human variants in other ISP marker genes, such as DUOX2, are associated with IBD^44,65,66^ and its deletion in mice promotes increased intestinal permeability^67^. LCN2, a biomarker of inflammation in multiple contexts^35^, is another ISP maker gene with nuanced functional roles. Deletion of LCN2 in mice in conjunction with IL10 deletion is associated with increased inflammation and colonic tumor formation^36,37^, whereas its deletion alone appears to be protective against dextran sodium sulfate-induced intestinal damage^68^. Since cells in the ISP state appear to exhibit expression of genes that could be either protective or detrimental, functional analyses including lineage-tracing and ablation studies are needed to truly understand their contribution to IBD.

The persistent presence of cells in an ISP state could contribute to improper orchestration of self-renewal and differentiation. RNA velocity analyses^69^ suggest that ISPs originate from stem and OLFM4^+^REG1A^+^ secretory progenitor cells but are predicted to become absorptive colonocytes (Supplemental Figure 6). These results are consistent with recent findings by Li et al^70^, indicating that a similar cell population termed “LND (LCN2, NOS2, DUOX2) cells” may differentiate from stem/progenitor cells to absorptive colonocytes in adult patients with Crohn’s disease. We analyzed raw data from Li et al. using our bioinformatic pipeline, and the converse (Li et al pipeline with our data), and found that cells in an ISP cell state are indeed the same population as LND cells (Supplemental figure 7A). Furthermore, just like a subset of “late” LND cells were assigned absorptive colonocyte lineage by Li et al, we also detect a small, isolated portion of ISPs nestled within the FABP1+ absorptive colonocyte cluster (Figure 1E-F, Supplemental Figure 7B). In addition to altered lineage commitment to absorptive enterocytes, at least some cells in an ISP state may represent a subpopulation of deep crypt secretory cells, which have been described in mice, however, we did not detect the marker genes^50,51,71^ for these cells in any distinct cluster within our data. Our finding that ISPs exhibit high fetal reprogramming gene expression further supports the notion that these cells exhibit marked plasticity, which may be associated with loss of lineage fidelity. Transient lineage plasticity is a hallmark of colonic regeneration, and emerging data suggest that founder progenitor cells expressing interferon targets can clonally expand to promote wound healing after wide-spread epithelial cell damage via dextran sodium sulfate (DSS)^72^. These findings, together with the presence of ISPs in healthy control subjects, suggest that the ISP state represents normal wound healing and that in IBD, the persistence of this state could contribute to IBD progression or even progression of colitis-associated colorectal cancer. In this vein, a recent study demonstrated that Paneth cells lacking the tumor suppressor gene *Apc* give rise to colonic tumors in the context of DSS-induced inflammation in mice^73^. Taken together, ISPs may be a part of typical wound healing process, however the present studies indicate that in IBD, there must be a mechanism by which ISPs persist.

We report that ISPs exhibit open chromatin and gene expression of ISP signature genes. ISP-specific genes are not expressed in intestinal stem cells despite their chromatin being accessible in Crohn’s disease versus healthy stem cells, demonstrating that ISP genes are epigenetically poised in stem cells and are transcriptionally activated in ISP cells in the presence of inflammatory stimuli. We confirmed this hypothesis using Crohn’s disease colonoids, which exhibit sustained ISP gene expression that can be robustly elicited via inflammatory stimuli. Furthermore, we identify the subset of ISP genes that are likely to drive this epigenetic memory via increased chromatin accessibility around pro-inflammatory transcription factor binding motifs. Recent studies in a mouse model of gastrointestinal acute graft-versus-host disease (GVHD) showed that *Lgr5*+ stem cells exhibited changes in oxidative phosphorylation genes in vivo that persisted in enteroid culture, including a functional reduction in oxygen consumption rates. In addition, post-GVHD stem cells exhibited reduced regenerative capacity based on organoid formation efficiency assays^74^. These changes were also associated with epigenetic memory, albeit via different signaling pathways than we describe. Reduced regenerative capacity of intestinal stem cells following in vivo challenge has also been demonstrated in mouse models of T cell-mediated tissue injury^75,76^. Together, studies in mice suggest that acute injury can yield epithelial changes that persist in organoid culture. However, whether epithelial changes observed in mouse models are sustained long term is not known. While our data in human organoids support the premise that the IBD epithelium exhibits memory likely due to prior inflammation, the duration of prior insult required to generate sustained epigenetic memory remains unclear. In addition, the extent to which the type or severity of injury dictates how the epigenome is rewired, is unknown.

Future studies are needed to define the relative contribution of DNA methylation, poised versus active enhancer activity, and other histone modifications to the emergence of the ISP state. While additional studies are needed to identify the ISPs as a therapeutic target, another clinical translation of this work is to define the extent to which abundance of cells in the ISP state can serve as a prognostic marker. This possibility could be especially important as an early marker in children with IBD to predict disease progression. In conclusion, our findings suggest that epigenetic memory of intestinal stem cells not only contributes to the persistence of a putatively pathogenic ISP cell state, but also provides new avenues for epithelial-targeted therapies and prognostic tools for IBD.

### Limitations

While our data are derived directly from patients, and while the sampled area was not actively inflamed, we cannot conclude that altered epigenetic states are not a result of continued exposure to systemic inflammatory signaling. However, the persistence of an ISP transcriptional signature in Crohn’s colonoids in vitro suggests that this state is, at least in part, an epithelial-intrinsic mechanism.

## METHODS

### Subject enrollment and demographics

This study was conducted with the approval of the Children’s Hospital of Philadelphia Institutional Review Board (IRB): #14-010826 (PI Dr. Judith Kelsen) and the University of Pennsylvania Perelman School of Medicine IRB: #814428 (PI Dr. Meenakshi Bewtra). For pediatric subjects, all parents of patients provided written informed consent. For adult subjects, all patients provided written informed consent. Biopsy specimens and histological samples were obtained from de-identified patients by Drs. Kelsen and Bewtra and their clinical colleagues. In total, we processed endoscopically non-inflamed biopsies from the ascending colon from 19 patients with Crohn’s disease and 23 control subjects. All pediatric patients were treatment naïve, with a range of disease activity (30% quiescent, 30% mild, 40% moderate disease). In the adult group, all of the patients were biologic-naïve, one patient was treatment-naïve, and 2 patients were not receiving treatment at the time of biopsy. Most adult patients were in remission (89%), and one patient (11%) had moderate disease. Detailed patient information, including age, sex, race and ethnicity, indications for endoscopy, is available in Supplemental Table 1.

### Sample processing and preparation for downstream analyses

Two biopsies per site were harvested in cryopreservation media for subsequent batch sample preparations for downstream analyses as published previously^34^. Briefly, samples were processed to separate epithelial-enriched and non-epithelial fractions which were processed separately for single cell RNA-seq, single nuclei ATAC-seq, and colonoid line generation. Single cell RNA-seq libraries were generated using Chromium Next GEM Single Cell 3’ GEM, Library & Gel Bead Kit v3.1 (10X GENOMICS INC) as per manufacturer’s instructions. Illumina NovaSeq 6000 was used to sequence libraries using parameters Read1: index5: index7:Read2:: 28:8:0:87 or 28:10:10:90.

### scRNA-seq QC and processing

We aligned and quantified the reads using 10X Cellranger 3.1.0 and built a gene expression matrix by combining all the available samples. Low quality cells expressing less than 1,000 genes and mitochondrial UMI (Unique Molecular Identifier) greater than 30% of the cell total were discarded. Only the genes that are expressed in more than 100 cells were used for analysis. The UMI counts for each gene were normalized by the cell totals followed by log transformation after addition of a pseudonumber^77^ to avoid negative and undefined log values.

Using the top 2000 genes with highest expression variation across the cells, we performed Principal Component Analysis (PCA) on the log normalized data with R preprocessCore package^77^. Top 10 dimensions of the PCA explaining the greatest variation in the data were used as input for non-linear dimensionality reduction with Uniform Manifold Approximation and Projection (UMAP)^78^. We ran clustering on the UMAP coordinates using density clustering function implemented in Monocle^79^. Finally, we assigned the cell types for clusters based on the expression patterns of the known cell markers curated from the literature. We verified that no cluster was disproportionately represented in select samples (Supplemental figure 1E).

### RNA velocity analysis

Spliced, unspliced and ambiguous expression matrices for each patient samples were generated with the tool velocyto.py v0.17.17. These matrices were then integrated into our previously generated Seurat (v4) object, which contains raw and integrated RNA expression profiles, as well as all the related meta information like annotated cell types. Only epithelial cells were extracted and included in the analysis. A UMAP was generated for this subset of cells based on 2000 highly variable genes across dataset. Each patient samples were then split from the integrated Seurat object and saved as H5ad format for downstream analysis. For each individual subsets, single cell RNA velocities and transcription rates were calculated using the spliced RNA oriented model as implemented in the package UniTVelo v0.2.5^80^. A transition matrix of epithelial clusters based on both RNA velocity and similarities among cells was calculated and combined using the methods implemented in CellRank v2.0.2^81^. The stream of velocities was plotted onto the embedding using the function VelocityKernel.plot_projection. Macrostates were inferred based on previously estimated transition matrix. Initial and terminal states were defined using the GPCCA-based workflow. The fate probabilities for all states were calculated for cells. The fate probabilities of all cells in the clusters were visualized using the function pl.aggregate_fate_probabilities with violin mode. Last, a triangular embedding plot was generated using the function pl.circular_projection.

### Clinical samples included in snATAC-seq analysis

snATAC libraries were generated from healthy and Crohn’s disease pediatric tissue samples of the ascending colon using Chromium Next GEM Single Cell ATAC Library & Gel Bead Kit v1.1 (10X GENOMICS INC) as per manufacturer’s instructions. Libraries were then sequenced on Illumina NovaSeq 6000 using parameters Read1: index5: index7:Read2::49:8:16:49. Samples from failed sequencing runs or with fewer than 100 high-quality sequenced cells were excluded from analysis.

### snATAC-seq QC and processing

Raw 10X scATAC-seq data were demultiplexed and aligned to the GRCh38 genome using the *mkfastq* and *count* functions from Cell Ranger ATAC (version 2.0.0). ArchR arrow files and a 500-bp bin-by-cell matrix were created for downstream QC and analysis with the ArchR package version 1.0.2. High-quality cell barcodes satisfying QC measures of 1. Number of fragments [2500-150,000] (Supplemental Figure 6A-B). 2. TSS enrichment score [>12] 3. Nucleosome ratio [0.5-10] 4. Blacklist ratio [<0.5] were retained. Doublets were inferred using *addDoubletScores() (k=10, knnMethod=”UMAP”)* and removed. Initial dimensionality reduction on bin features was performed using iterative latent sematic indexing (LSI) implemented by *addIterativeLSI() (iterations=4, varFeatures=25000, resolution=0.2)*. Harmony was applied to the IterativeLSI dimensionality reduction to correct for batch effects across patient samples. Cells were clustered in the harmonized space using *addClusters() (method=”Seurat”, resolution=1.7)* and results were visualized using UMAP. Gene scores were computed as a function of chromatin accessibility within the gene body and putative upstream distal regulatory elements using *addImputeWeights().* Accessible peaks were called using ArchR’s implementation of macs2 on pseudobulk replicates. A peak-by-cell matrix (insertions per peak and cell) was generated and used as input to ChromVAR, which calculates bias-corrected enrichment of TF motif accessibility on a per-cell basis.

### snATAC-seq cell type annotation

Broad cell annotations were manually obtained by examining clusters for gene activity enrichment of canonical lineage-specific marker genes and gene signatures from Smillie et al.^9^ for Epithelial, Immune, and Stromal compartments. Epithelial compartment cells were subsetted and re-processed to identify finer epithelial cell types. Epithelial cell annotations were manually obtained by examining clusters for gene activity enrichment of canonical cell type marker genes, Burclaff et al. 2022 epithelial signatures, and custom epithelial signatures.

### Heatmap and box plot analysis

Heatmap showing the relative Gene Activity (z-score) of intestinal epithelial gene signatures (rows) across the identified epithelial cell types were plotted using hierarchical clustering pheatmap v1.0.12 function in R.

Published fetal and regenerative signatures in epithelial sub-clusters were aggregated based on individual patients. Average expression of these aggregated signatures were calculated based on each epithelial cell types, and then visualized via box plots separated by different conditions using the function geom_half_boxplot implemented in gghalves v0.1.4.

Peak conversion values for ISP marker genes HLA-DPA1, HLA-DRA, and CD74, in grouped progenitor cells (stem cells, TA, early progenitor clusters), were visualized via box plots compared from control subjects versus Crohn’s disease patients, using the function geom_boxplot implemented in ggplot2 package.

### Colonoid line generation, organoid formation assays, inflammatory cocktail stimulation

Roughly 1/3 of the epithelial single cell suspension prepared for sequencing was used to establish colonoid cultures as described^82^. Briefly, 4,500 live cells from dissociated crypts were seeded in a 50 uL dome of 80% Matrigel, allowed to solidify for 45 minutes at 37°C. IntestiCult OGM (StemCell Tech #06010) was supplemented with ROCK inhibitor Y27632 (working concentration 10 uM) at seeding, and Primocin (InvivoGen #ant-pm-05) was used in the first passage to prevent contamination. After 14 days of culture, the colonoids were dissociated via trypsinization and seeded (1) for expansion at 2,000-5,000 live cells per 50 uL dome of 80% Matrigel or (2) for organoid formation efficiency (OFE) assay at 500 live cell in 10 uL domes of 80% Matrigel. The cultures were imaged on days 7 and 14 post-seeding using Keyence BZX microscope. All experiments were carried out in differentiation medium^83^ (50% L-WRN conditioned medium^84^, 1xB27 supplement, 50 ng/mL FGF-2, 100 ng/mL IGF-1, 10 nM Gastrin, 1 mM N-Acetylcysteine, 500nM A83-01). Inflammatory cocktail treatment was carried out by adding recombinant human TNFalpha (Peprotech #300-01A, final concentration 100ng/mL), IL-1beta (Peprotech # 200-01B, final concentration 20 ng/mL), IL-6 (Peprotech # 200-06, final concentration 10 ng/mL), and *S. typhimurim* flagellin (InvivoGen # FLA-ST, final concentration 1 ug/mL) for 24 h.

### Macrophage differentiation from monocytes

For each experiment, 10x10^6^ monocytes from healthy human donors were acquired from the Human Immunology Core at the University of Pennsylvania (RRID:SCR_022380) for differentiation into macrophage following the protocol established by STEMCELL Technologies. Briefly, monocytes were seeded into a 12 well plate, at 1x10^6^ monocytes/well, and allowed to differentiate for 6 days in ImmunoCult-SF (STEMCELL Technologies: Cat# 10961) supplemented with recombinant human M-CSF (STEMCELL Technologies: Cat# 78057; 50ng/mL). On day 4 of culture, the macrophages were activated with lipopolysaccharide (LPS; 10ng/mL) and IFNgamma (50ng/mL) and on the 6th day, were harvested for co-culturing with colonoids. For quality control, 1x10^6^ monocytes were cultured separately to access the percentage of macrophages (CD14^+^CD64^+^CD80^+^) that were derived from the monocytes (CD14^+^CD64^-^CD80^-^) using flow cytometry analysis.

### Colonoid-macrophage co-culture

Following macrophage activation, on the 6th day of culture, the macrophages were harvested for co-culturing with colonoids. In short, the ImmunoCult-SF Macrophage medium was removed and Accutase (STEMCELL Technologies: Cat# 07920) was added to the macrophages which were then incubated at 37°C for 15 minutes, then the Accutase was inactivated by 2x volume of 0.5% BSA in phosphate buffered saline (PBS). The cell suspension was transferred to a conical tube, centrifuged for 5 minutes at 300g and resuspended in medium containing a ratio 1:1 of 1 ImmunoCult-SF to enteroid differentiation medium (EDM). Colonoids were harvested on day 12-13 post-seeding, washed with ice-cold PBS, and re-embedded in 100ul of Matrigel containing 3x10^5^ macrophages and ∼100 colonoids per dome. The lines were co-cultured for another 24 hours in media that was a 1:1 ratio of ImmunoCult-SF and EDM containing flagellin (1ug/mL).

### Single cell dissociation, FACS sorting and flow cytometry analysis of colonoids

Colonoids (untreated, cytokine-treated, co-cultured) were harvested in PBS and centrifuged at 700g for 3 minutes. The samples were resuspended in TrypLE Express (Gibco: Cat# 12605-010) containing DNASE I (Sigma Aldrich: Cat# 10104159001; 0.50U/ml) and incubated in a thermomixer at 37C rotating at 800rpm for 12 minutes to obtain a single cell suspension. The TrypLE was inactivated in sorting buffer (2%BSA in 1x Hanks Balanced Salt Solution supplemented with 25mM HEPES, 0.5U/mL DNASEI and 10uM Y-27632HCl), and the samples subjected to extracellular staining for 30 minutes at 4C in the dark. For FACS sorting experiments, samples were sorted into TRI-Reagent (Sigma Aldrich: Cat# T3934-100ML) using either the Cytek Aurora or the FACS Melody. For flow cytometry analysis, the samples were fixed in Cytofix (BD Biosciences: Cat# 554655) for 15-20 minutes prior to analysis on the BD LSR Fortessa.

| Antibody | Conjugation | Manufacturer | Catalog | Dilution |
| --- | --- | --- | --- | --- |
| LIVE/DEAD<br>fixable blue dead<br>cell stain kit | N/A | Invitrogen | L34961A | 1 to 1000 |
| DAPI | N/A | Sigma Aldrich | D9542 | 1:10,000, from<br>stock of 1mg/ml |
| HLA-DP | PE | BD<br>Pharmingen | 566825 | 1 to 20 |
| HLA-DR | PerCP/Cy5.5 | Biolegend | 327020 | 1 to 20 |
| EpCAM | FITC | Biolegend | 324204 | 1 to 20 |
| CD45 | APC/Fire750 | Biolegend | 368517 | 1 to 20 |
| CD64 | APC | Biolegend | 305013 | 1 to 20 |
| CD14 | PerCP/Cy5.5 | Biolegend | 325621 | 1 to 20 |
| CD80 | FITC | Biolegend | 305205 | 1 to 20 |

### Quantification of ISP induction in macrophage-colonoid co-cultures

Since the degree of macrophage differentiation varied dramatically based on the donor (Supplemental figure 5D), when we plotted the %ISPs among the total epithelium in co-culture experiments and saw variability in degree of ISP induction (1.6%-12.9% in control colonoids vs 5.1% to 38.9% in Crohn’s colonoids, Supplemental Figure 5E), we opted to normalize % ISPs to the degree of macrophage differentiation efficiency (Supplemental figure 5D).

### Colonoid FFPE Embedding & Histology

Colonoids were harvested by dislodging the Matrigel with ice-cold DPBS and fixed at 4°C using 4% paraformaldehyde for an hour, followed by a wash with DPBS. After pelleting, fixed colonoids were resuspended in 80 uL of warmed embedding gel (2% bacto-agar + 2.5% gelatin) and transferred to chilled parafilm-wrapped block. Droplets of colonoids embedded in the gel were allowed to solidify at 4°C and then each placed into a histology cassette and stored in 70% EtOH for processing. Embedding into paraffin blocks, sectioning, and Hematoxylin and eosin staining were carried out by Center for Molecular Studies in Digestive and Liver Diseases and the Molecular Pathology and Imaging Core (RRID:SCR_022420).

### Immunofluorescence

FFPE slides were deparaffinized and rehydrated. Antigen retrieval was done in a pressure cooker (Antigen Retriever 2100) using Tris-EDTA antigen retrieval buffer (10 mM Tris base, 1 mM EDTA solution, 0.05% Tween 20, pH 9.0). Sections were permeabilized with PBS-T (1X DPBS with 0.1% Triton-X) for 5 minutes and then washed. All washes in between steps were done with 1X DPBS. Slides were blocked at room temperature for an hour (10% donkey serum and 1% BSA in DPBS). Primary antibodies were diluted in 1% BSA in DPBS and incubated overnight at 4°C. After washing, slides were incubated with secondary antibodies and DAPI (1:5000 dilution of 1mg/mL stock) for 30 minutes in the dark at room temperature. Slides were washed and then mounted using Prolong™ Gold Antifade Mountant (P36930, ThermoFisher Scientific). Slides were imaged on a Leica Stellaris 5 Laser-scanning confocal microscope at the Cell and Developmental Biology Microscope Core (RRID SCR_022373).

| Primary Antibody | Manufacturer | Catalog | Dilution |
| --- | --- | --- | --- |
| Rabbit Recombinant Monoclonal anti-HLA-DR [EPR3692] | Abcam | ab92511 | 1:100 |
| Purified Mouse anti-human CD74 Antibody | BioLegend | 326802 | 1:100 |
| Goat Polyclonal anti-LCN2 | Proteintech | AF1757 | 1:40 |
| Purified Mouse anti-ECAD | Cell Signaling | 610182 | 1:50 |
| Secondary Antibody |  | Catalog |  |
| Cy™2 AffiniPure™ Donkey Anti-Rabbit IgG | Jackson IR | 711-225-152 | 1:600 |
| Cy3 AffiniPure™ Donkey Anti-Mouse IgG | Jackson IR | 715-165-151 | 1:600 |
| Cy™5 AffiniPure™ Donkey Anti-Goat IgG | Jackson IR | 705-175-147 | 1:600 |

### Image Processing, Analysis & Quantification

Images acquired on the Leica Stellaris 5 were processed using ImageJ Fiji v2.14.0/1.54f. Z-stacks were compressed with maximum intensity projection to generate tiff files. For patient biopsies, the following contrast/brightness settings were applied in 8-bit range (0-255) to each image: Channel 1 (DAPI, blue) min: 15 max: 125; Channel 2 (HLA-DR, cyan) min: 15 max: 150; Channel 3 (CD74, yellow) min: 15 max: 175; Channel 4 (LCN2, red) min: 15 max: 255. Scale bar was set to 100 microns. For inset images of patient biopsies, the following contrast/brightness settings were applied in 8-bit range (0-255) to each image: Channel 1 (DAPI, blue) min: 15 max: 100; Channel 2 (HLA-DR, cyan) min: 15 max: 124; Channel 3 (CD74, yellow) min: 15 max: 175; Channel 4 (LCN2, red) min: 15 max: 150. Scale bar was set to 25 microns. Where applicable, contrast/brightness settings for ECAD (Channel 3, green) were set in 8-bit range to min: 10 max: 110. For organoids, the following contrast/brightness settings were applied in 8-bit range (0-255) to each image: Channel 1 (DAPI, blue) min: 0-5 max: 25-35; Channel 2 (HLA-DR, cyan) min: 15 max: 50; Channel 3 (CD74, yellow) min: 0 max: 65; Channel 4 (LCN2, red) min: 15 max: 100.To quantify ISPs in patient biopsies, images were imported into QuPath v0.5.0 and a representative image (biopsy from patient with Crohn’s Disease) was selected for setting quantification threshold and counts (see QuPath Script). Before running the script with newly defined “ISP” classifier, crypts were outlined as annotations in each image using ECAD-stained images for reference and to distinguish epithelial from non-epithelial cells. Therefore, the classifier was applied only to crypts across all images and quantification and further analysis was computed by crypt (“ISPs per crypt”). To quantify ISPs in organoids, images were also imported into QuPath and single cells with all 3 markers (HLA-DR/CD74/LCN2) were annotated. These annotations were exported as counts to quantify number of ISPs per image.

### RNA Isolation & qPCR

Colonoids were harvested and resuspended in TRIzol™ Reagent (Catalog #15596026, Ambion). RNA was extracted using a Direct-zol RNA Microprep Kit (Catalog #R2062, Zymo Research). cDNA was generated using the High-Capacity cDNA Reverse Transcription Kit (Catalog #4368814, Invitrogen). Real-time quantitative polymerase chain reaction was performed with TaqMan™ probes listed below and TaqMan™ Fast Universal PCR Master Mix 2X (Catalog #4352042).

| Taqman Probe | Assay ID |
| --- | --- |
| TBP1 | Hs00427621_m1 |
| LCN2 | Hs01008571_m1 |
| HLA-DRA | Hs00219575_m1 |
| HLA-DPA1 | Hs00410276_m1 |
| CD74 | Hs00269961_m1 |

## ACKNOWLEDGEMENTS

We are grateful to the participating patients and families, as well as nurses and staff at the CHOP and Penn gastrointestinal endoscopic suites for supporting our research. We also thank Dr. Sebastian Pott (University of Chicago), Dr. Lori Coburn (Vanderbilt University), and members of the Helmsley Gut Cell Atlas consortium for insightful conversations. We thank the following scientific cores and centers: the Center for Molecular Studies in Digestive and Liver Diseases (P30DK050306) and the Molecular Pathology and Imaging Core (RRID: SCR_022420), the University of Pennsylvania Cell and Developmental Biology Microscope Core (RRID SCR_022373), the Flow Cytometry Core at the Children’s Hospital of Philadelphia, CHOP Gastrointestinal Epithelium Modeling Program. This publication is part of the Gut Cell Atlas Crohn’s Disease Consortium funded by the Leona M. and Harry B. Helmsley Charitable Trust and is supported by a grant from Helmsley to the Children’s Hospital of Philadelphia. www.helmsleytrust.org/gut-cell-atlas/. The study was also supported by funding from Children’s Hospital of Philadelphia institutional development funds (KEH).

---

**Supplemental Figure 1 (related to Figure 1):**
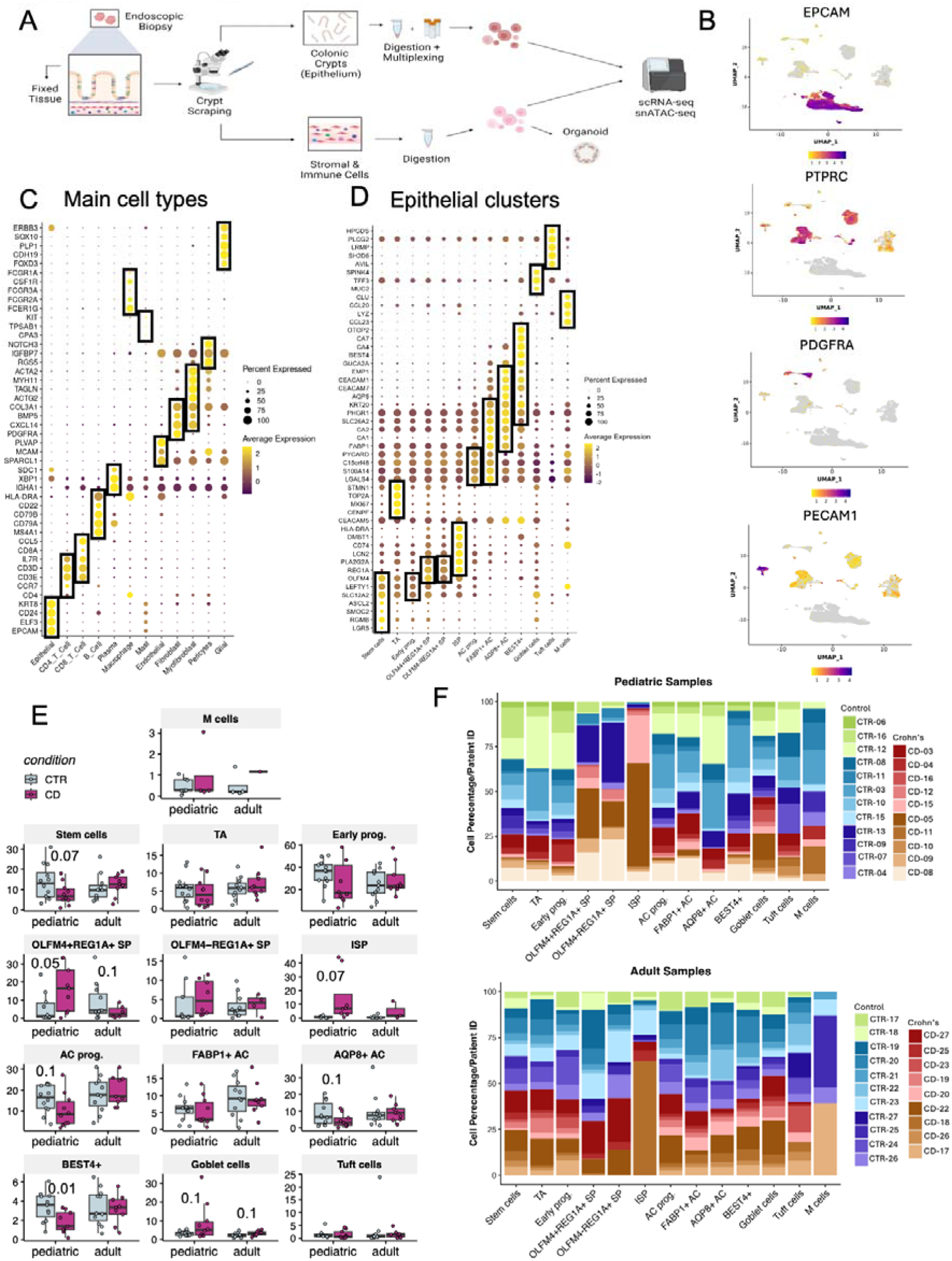
Single cell composition of colonic biopsies from healthy controls and Crohn’s disease patients. (A) Study schematic: endoscopic biopsies were cryopreserved upon collection. For generation of scRNA-seq and snATAC-seq libraries, epithelium-enriched (via crypt scraping) and lamina propria fractions were processed separately. (B) Expression of key marker genes for the epithelial (EPCAM), immune (PTPRC), fibroblast (PDGFRA), endothelial (PECAM1) lineages plotted on the UMAP. (C and D) Differentially expressed genes defining the cell clusters (C) and epithelial cell sub-clusters (D). The size of the dot represents the fraction of cells within cluster expressing the gene and color represents expression level. (E) Epithelial sub-cluster abundance, per individual subject (P≤ 0.1 shown over respective comparisons). (F) Contribution of individual subjects to each epithelial sub-cluster.

**Supplemental figure 2 (Related to figure 2):**
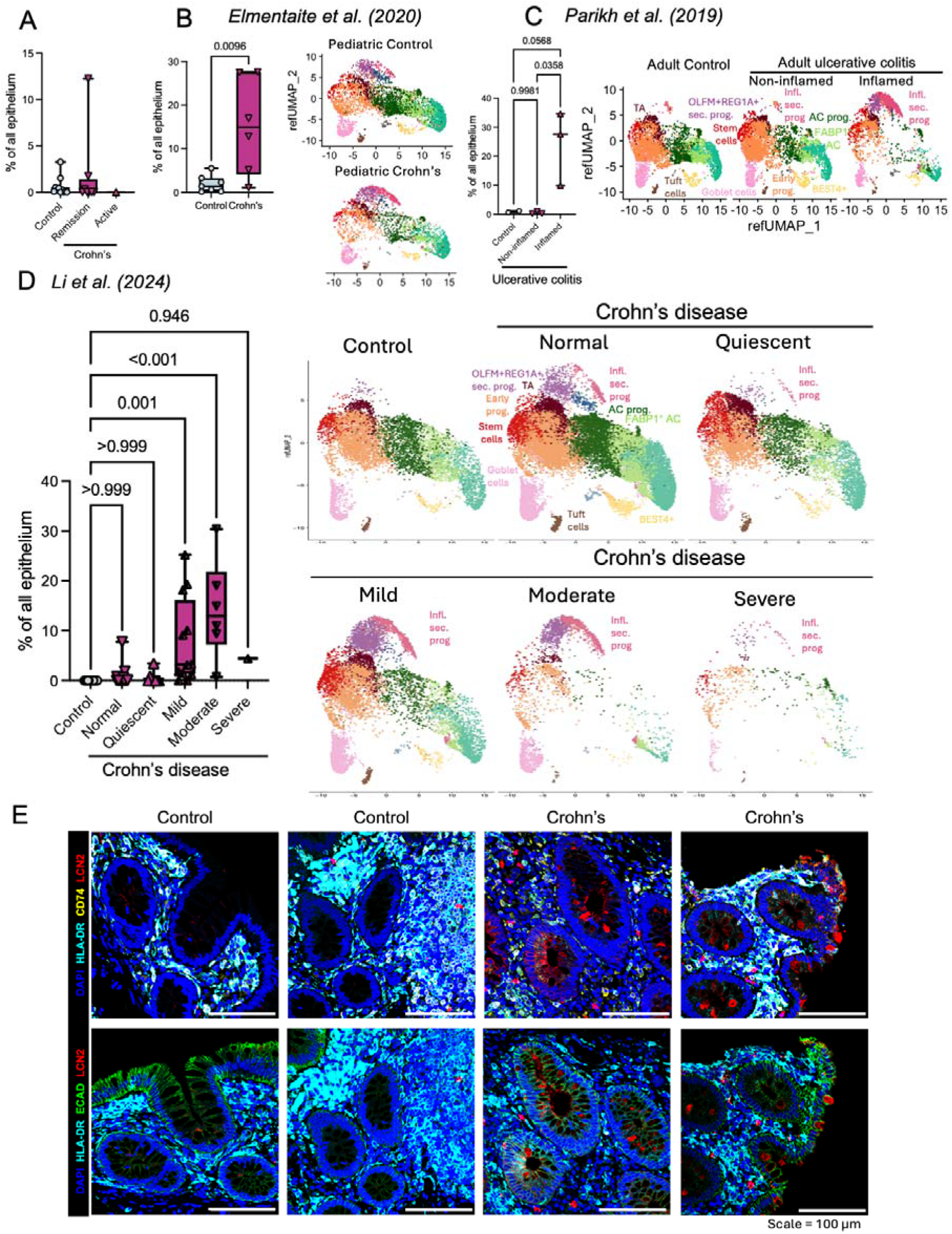
Inflammatory secretory progenitors constitute a cell state specific to Crohn’s disease. (A) Relative abundance of the ISP cluster in total epithelium for individual adult subjects. (B) UMAP projections on a published dataset (E-MTAB-8901; pediatric Crohn’s disease). (C) UMAP projections on a published dataset (GSE116222; adult ulcerative colitis). (D) UMAP projections on a published dataset (GSE266546; adult Crohn’s disease). Crohn’s disease activity classification presented as reported in the original study. (E) Additional replicates of immunostaining for ISP markers in colonic biopsies from study subjects.

**Supplemental Figure 3 (related to Figure 3).**
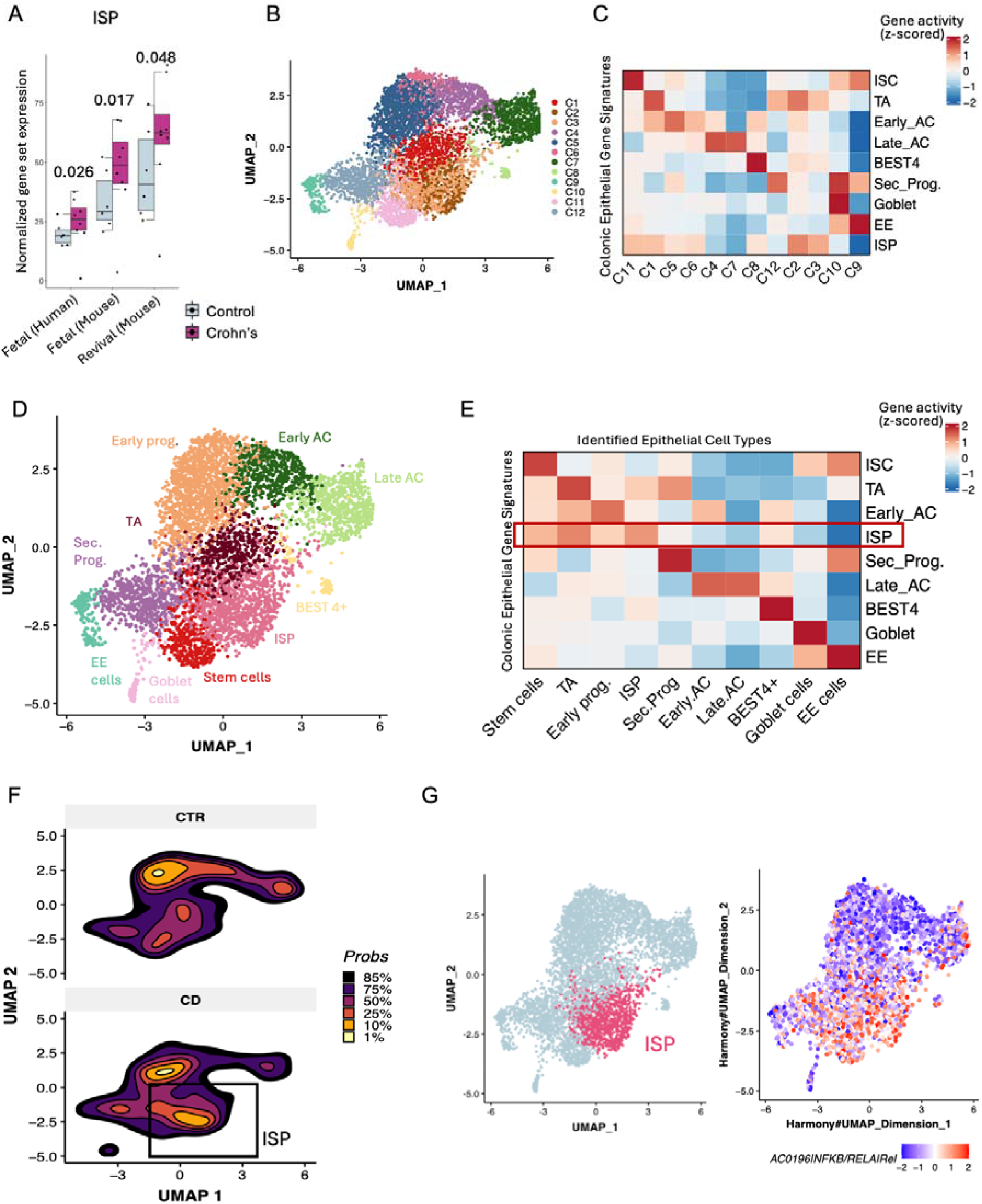
(A) Relative abundance of the ISP cluster in total epithelium for individual pediatric subjects (in active Crohn’s disease, biopsies were taken from areas adjacent to inflammation), P values (ordinary one-way ANOVA) presented on the plots. (B) UMAP representation of all epithelial cells sequenced from pediatric subjects color-coded by cell cluster. A total of 12 district clusters were obtained. (C) Heatmap showing the relative Gene Activity (z-score) of intestinal epithelial gene signatures (rows) across the identified epithelial cell clusters (columns). (D) UMAP representation of all epithelial cells sequenced from pediatric subjects annotated and color-coded by cell type. (E) Heatmap showing the relative Gene Activity (z-score) of intestinal epithelial gene signatures (rows) across the identified epithelial cell types (columns). (F) Density map showing UMAP embedding of chromatin accessibility features for Crohn’s and control samples. Density contours represent regions of highest cell density from each condition with color gradient reflecting the location of highest cell density. (G) UMAP of all epithelial cells from pediatric samples with ISP cells highlighted in pink (left). UMAP showing the enrichment of NFkB motifs across cell clusters (right).

**Supplemental Figure 4 (related to Figure 4):**
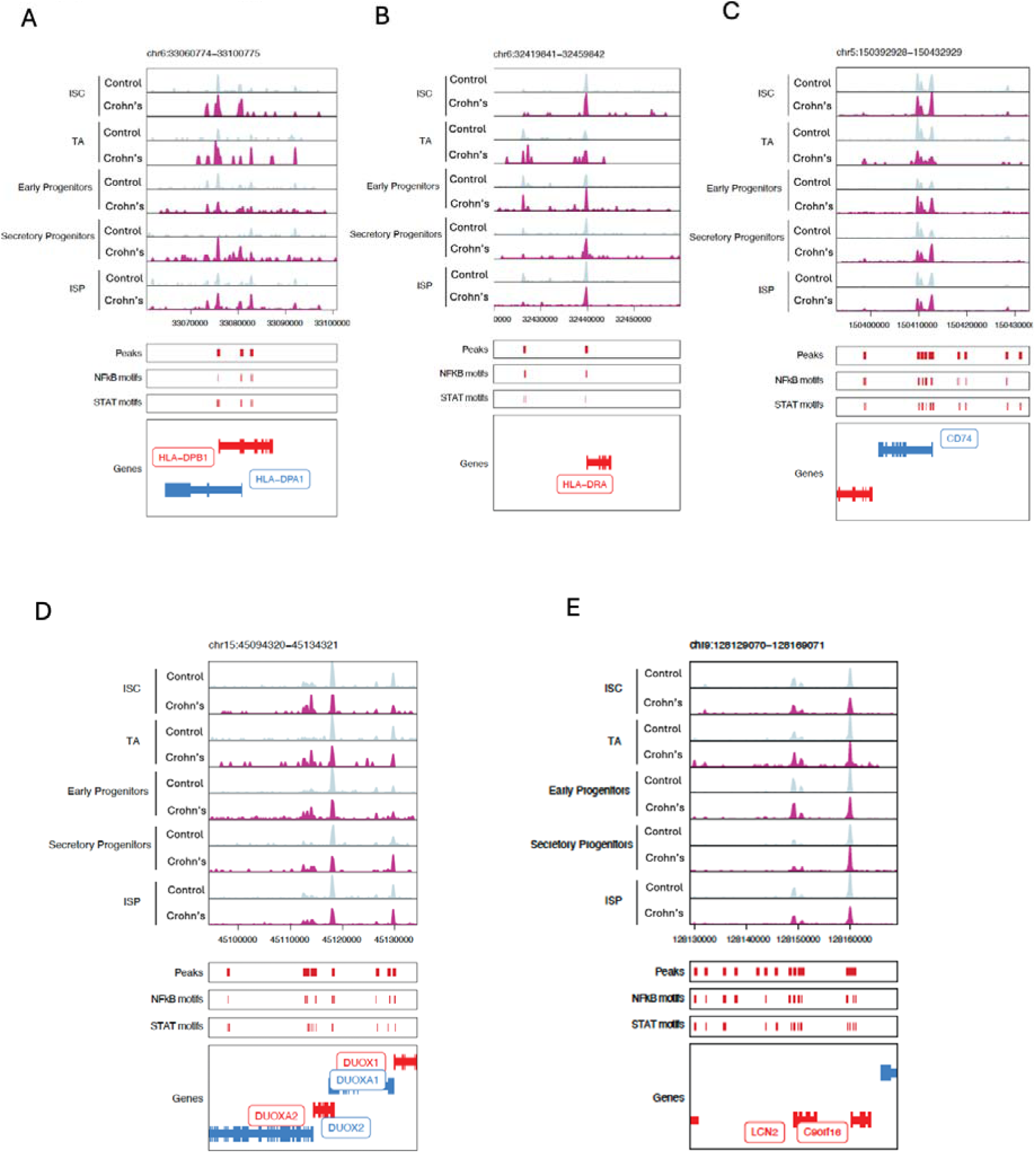
Select ISP signature genes contain open chromatin peaks enriched in Crohn’s disease, in multiple epithelial sub-clusters. Open chromatin peaks in stem cell, TA, early progenitors, secretory progenitors, ISP, clusters, with traces from control and Crohn’s disease subjects separated by color. Corresponding gene maps positioned below the chromatin traces.

**Supplemental Figure 5 (related to Figure 5):**
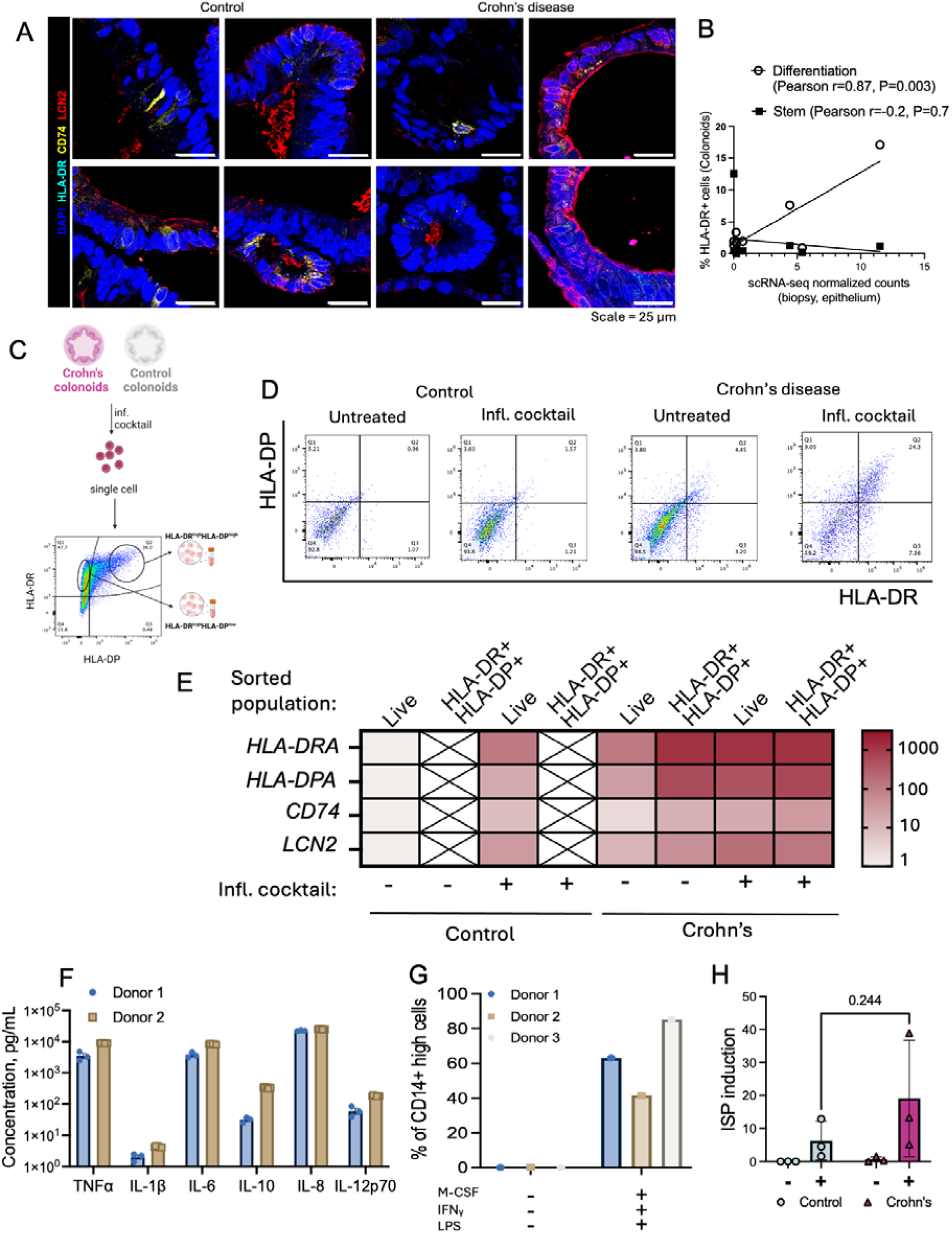
An inflammatory cocktail stimulates ISP state *in vitro*. (A) Single channel images related to Figure 5B. (B) Correlation analysis of HLA-DRA gene expression in the study subjects’ biopsies (scRNA-seq data, epithelial cluster) vs HLA-DR protein expression in colonoids from corresponding subjects (flow cytometry), comparing colonoids grown either in stem cell-enriching expansion medium (black squares) or differentiation medium (open circles). (C) Schematic: Colonoids from from control subject or Crohn’s disease patient were treated with the inflammatory cocktail for 24h and subjected to flow cytometry sorting of HLA-DR+ and HLA-DR+HLA-DP+ populations, as well as total live cells to use as control. (D) Representative flow cytometry plots from the sorting experiment described. Cell frequencies for the double-negative, HLA-DR+, HLA-DP+, and double-positive populations are presented on the plots. (E) Expression of ISP marker genes HLA-DPA1, HLA-DRA, CD74, and LCN2, in cell populations sorted as described in (C) from control or Crohn’s colonoids. One line each from control subject or Crohn’s disease patient, N=3 independent experiments repeated in different passages (P=4-6), representative data from one experiment are shown. (F) Quantification of secretion of the cytokines TNFa, IL-1b, IL-6, IL-10, IL-8, IL-12p70 by macrophages differentiated and activated ex vivo prior to co-embedding with colonoids. (G) Assessment of macrophage activation via flow cytometry for co-expression of CD64 and CD80 within the CD14high monocyte population. N=3 independent donors. (H) Quantification of % EpCAM+HLA-DR+HLA-DP+ cells without normalization to the degree of macrophage activation and polarization, in colonoid mono-cultures or co-cultures with M1 macrophages.

**Supplemental figure 6:**
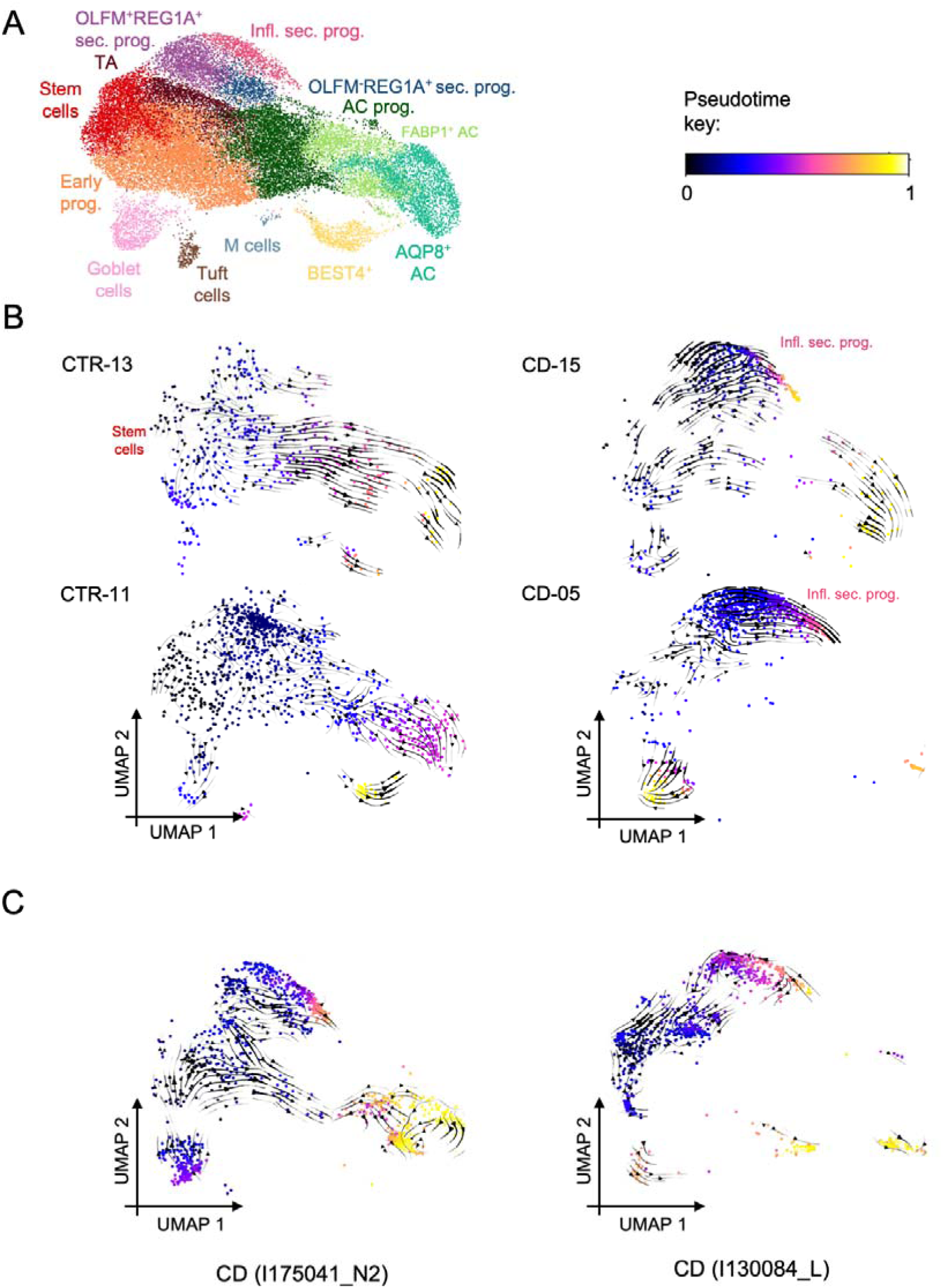
ISP cells contribute to the absorptive lineage. (A) Epithelial cluster UMAP key and Pseudotime color legend for plots in (B-C). (B) Predicted future transcriptional states for individual cells plotted on epithelial UMAPs from representative pediatric subjects. (B) Predicted future transcriptional states for individual cells plotted on the epithelial UMAP. Two samples of adult Crohn’s disease from Kong et al. dataset (DUOS-000145, DUOS-000146) are shown.

**Supplemental figure 7:**
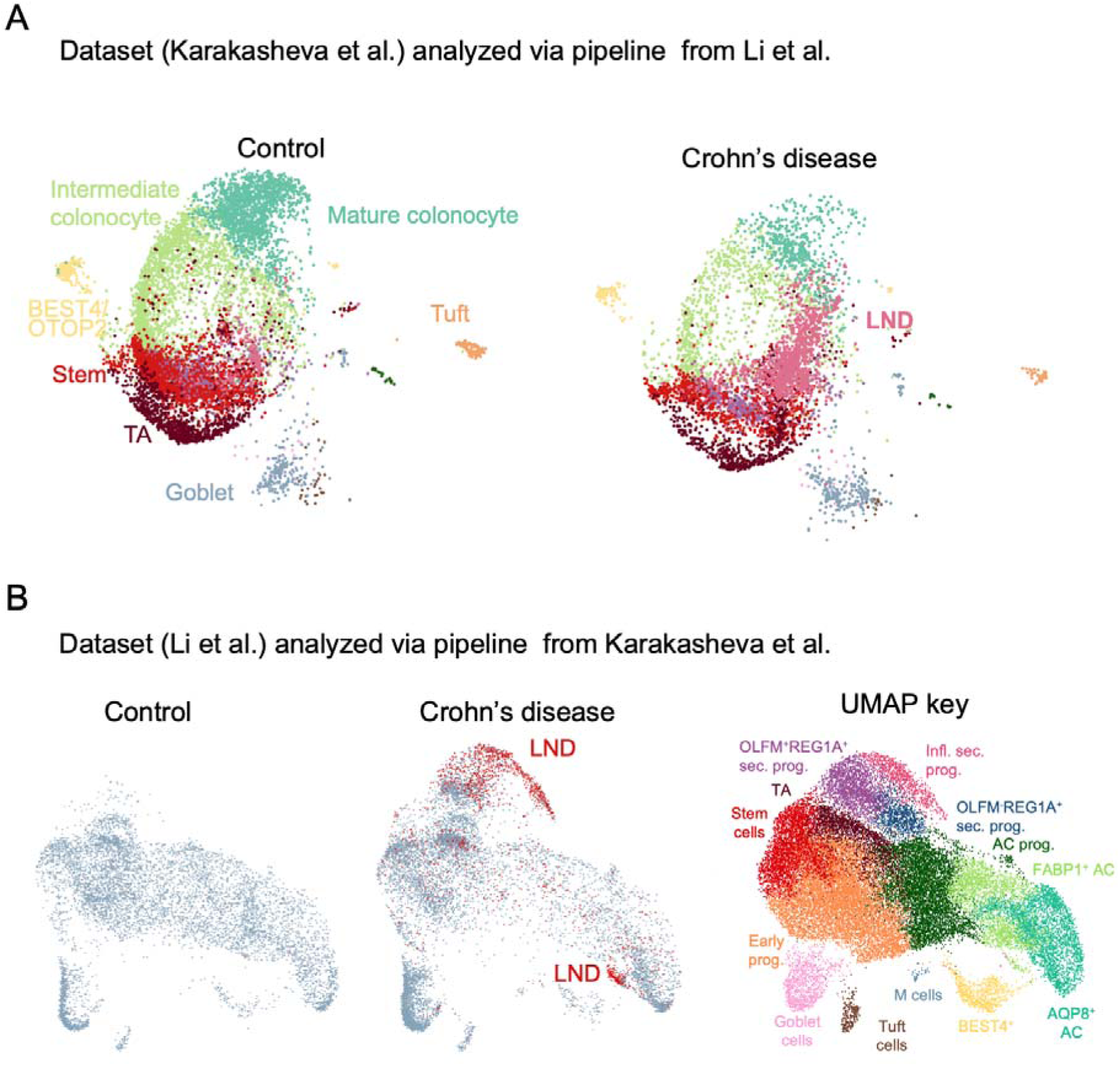
Cells in ISP state are synonymous with recently described LND cells. (A) Epithelial transcriptomes from this study were analyzed through the pipeline from Li et al. The cells identified as LNDs match their position on the UMAP in original paper. (B) Epithelial transcriptomes (colon) from Li et al. were analyzed through the pipeline from this study, and the LND cells are highlighted in red, demonstrating that the cluster is split between the secretory and absorptive lineages. The color key to position of epithelial clusters is provided on the left.

## REFERENCES

1. Coskun, M. (2014). Intestinal epithelium in inflammatory bowel disease. Front Med (Lausanne) 1, 24. 10.3389/fmed.2014.00024.

2. Villablanca, E.J., Selin, K., and Hedin, C.R.H. (2022). Mechanisms of mucosal healing: treating inflammatory bowel disease without immunosuppression? Nature Reviews Gastroenterology & Hepatology 2022 19:8 19, 493–507. 10.1038/s41575-022-00604-y.

3. Otte, M.L., Lama Tamang, R., Papapanagiotou, J., Ahmad, R., Dhawan, P., and Singh, A.B. (2023). Mucosal healing and inflammatory bowel disease: Therapeutic implications and new targets. World J Gastroenterol 29, 1157–1172. 10.3748/wjg.v29.i7.1157.

4. Molodecky, N.A., Soon, I.S., Rabi, D.M., Ghali, W.A., Ferris, M., Chernoff, G., Benchimol, E.I., Panaccione, R., Ghosh, S., Barkema, H.W., et al. (2012). Increasing incidence and prevalence of the inflammatory bowel diseases with time, based on systematic review. Gastroenterology 142, 46–54.e42; quiz e30. 10.1053/j.gastro.2011.10.001.

5. Rosen, M.J., Dhawan, A., and Saeed, S.A. (2015). Inflammatory Bowel Disease in Children and Adolescents. JAMA Pediatr 169, 1053–1060. 10.1001/jamapediatrics.2015.1982.

6. Ruemmele, F.M., El Khoury, M.G., Talbotec, C., Maurage, C., Mougenot, J.-F., Schmitz, J., and Goulet, O. (2006). Characteristics of Inflammatory Bowel Disease With Onset During the First Year of Life. Journal of Pediatric Gastroenterology and Nutrition 43, 603–609. 10.1097/01.mpg.0000237938.12674.e3.

7. Mitsialis, V., Wall, S., Liu, P., Ordovas-Montanes, J., Parmet, T., Vukovic, M., Spencer, D., Field, M., McCourt, C., Toothaker, J., et al. (2020). Single-Cell Analyses of Colon and Blood Reveal Distinct Immune Cell Signatures of Ulcerative Colitis and Crohn’s Disease. Gastroenterology 159, 591–608.e10. 10.1053/j.gastro.2020.04.074.

8. Parikh, K., Antanaviciute, A., Fawkner-Corbett, D., Jagielowicz, M., Aulicino, A., Lagerholm, C., Davis, S., Kinchen, J., Chen, H.H., Alham, N.K., et al. (2019). Colonic epithelial cell diversity in health and inflammatory bowel disease. Nature 567, 49–55. 10.1038/s41586-019-0992-y.

9. Smillie, C.S., Biton, M., Ordovas-Montanes, J., Sullivan, K.M., Burgin, G., Graham, D.B., Herbst, R.H., Rogel, N., Slyper, M., Waldman, J., et al. (2019). Intra-and Inter-cellular Rewiring of the Human Colon during Ulcerative Colitis. Cell 178, 714–730.e22. 10.1016/j.cell.2019.06.029.

10. Kinchen, J., Chen, H.H., Parikh, K., Antanaviciute, A., Jagielowicz, M., Fawkner-Corbett, D., Ashley, N., Cubitt, L., Mellado-Gomez, E., Attar, M., et al. (2018). Structural Remodeling of the Human Colonic Mesenchyme in Inflammatory Bowel Disease. Cell 175, 372–386.e17. 10.1016/j.cell.2018.08.067.

11. Elmentaite, R., Ross, A.D.B., Roberts, K., James, K.R., Ortmann, D., Gomes, T., Nayak, K., Tuck, L., Pritchard, S., Bayraktar, O.A., et al. (2020). Single-Cell Sequencing of Developing Human Gut Reveals Transcriptional Links to Childhood Crohn’s Disease. Developmental Cell 55, 771–783.e5. 10.1016/j.devcel.2020.11.010.

12. Kong, L., Pokatayev, V., Lefkovith, A., Carter, G.T., Creasey, E.A., Krishna, C., Subramanian, S., Kochar, B., Ashenberg, O., Lau, H., et al. (2023). The landscape of immune dysregulation in Crohn’s disease revealed through single-cell transcriptomic profiling in the ileum and colon. Immunity 0. 10.1016/J.IMMUNI.2023.01.002.

13. Yui, S., Azzolin, L., Maimets, M., Pedersen, M.T., Fordham, R.P., Hansen, S.L., Larsen, H.L., Guiu, J., Alves, M.R.P., Rundsten, C.F., et al. (2018). YAP/TAZ-Dependent Reprogramming of Colonic Epithelium Links ECM Remodeling to Tissue Regeneration. Cell Stem Cell 22, 35–49.e7. 10.1016/j.stem.2017.11.001.

14. Chen, L., Qiu, X., Dupre, A., Pellon-Cardenas, O., Fan, X., Xu, X., Rout, P., Walton, K.D., Burclaff, J., Zhang, R., et al. (2023). TGFB1 induces fetal reprogramming and enhances intestinal regeneration. Cell Stem Cell 30, 1520–1537.e8. 10.1016/j.stem.2023.09.015.

15. Nusse, Y.M., Savage, A.K., Marangoni, P., Rosendahl-Huber, A.K.M., Landman, T.A., de Sauvage, F.J., Locksley, R.M., and Klein, O.D. (2018). Parasitic helminths induce fetal-like reversion in the intestinal stem cell niche. Nature 559, 109–113. 10.1038/s41586-018-0257-1.

16. Mustata, R.C., Vasile, G., Fernandez-Vallone, V., Strollo, S., Lefort, A., Libert, F., Monteyne, D., Pérez-Morga, D., Vassart, G., and Garcia, M.-I. (2013). Identification of Lgr5-independent spheroid-generating progenitors of the mouse fetal intestinal epithelium. Cell Rep 5, 421–432. 10.1016/j.celrep.2013.09.005.

17. Fernandez Vallone, V., Leprovots, M., Strollo, S., Vasile, G., Lefort, A., Libert, F., Vassart, G., and Garcia, M.I. (2016). Trop2 marks transient gastric fetal epithelium and adult regenerating cells after epithelial damage. Development 143, 1452–1463. 10.1242/dev.131490.

18. Wang, Y., Chiang, I.L., Ohara, T.E., Fujii, S., Cheng, J., Muegge, B.D., Ver Heul, A., Han, N.D., Lu, Q., Xiong, S., et al. (2019). Long-Term Culture Captures Injury-Repair Cycles of Colonic Stem Cells. Cell 179, 1144. 10.1016/J.CELL.2019.10.015.

19. Sheaffer, K.L., Kim, R., Aoki, R., Elliott, E.N., Schug, J., Burger, L., Schübeler, D., and Kaestner, K.H. (2014). DNA methylation is required for the control of stem cell differentiation in the small intestine. Genes Dev 28, 652–664. 10.1101/gad.230318.113.

20. Raab, J.R., Tulasi, D.Y., Wager, K.E., Morowitz, J.M., Magness, S.T., and Gracz, A.D. (2020). Quantitative classification of chromatin dynamics reveals regulators of intestinal stem cell differentiation. Development 147, dev181966. 10.1242/dev.181966.

21. Del Poggetto, E., Ho, I.-L., Balestrieri, C., Yen, E.-Y., Zhang, S., Citron, F., Shah, R., Corti, D., Diaferia, G.R., Li, C.-Y., et al. (2021). Epithelial memory of inflammation limits tissue damage while promoting pancreatic tumorigenesis. Science 373, eabj0486. 10.1126/science.abj0486.

22. Falvo, D.J., Grimont, A., Zumbo, P., Fall, W.B., Yang, J.L., Osterhoudt, A., Pan, G., Rendeiro, A.F., Meng, Y., Wilkinson, J.E., et al. (2023). A reversible epigenetic memory of inflammatory injury controls lineage plasticity and tumor initiation in the mouse pancreas. Dev Cell 58, 2959–2973.e7. 10.1016/j.devcel.2023.11.008.

23. Lim, A.I., McFadden, T., Link, V.M., Han, S.-J., Karlsson, R.-M., Stacy, A., Farley, T.K., Lima-Junior, D.S., Harrison, O.J., Desai, J.V., et al. (2021). Prenatal maternal infection promotes tissue-specific immunity and inflammation in offspring. Science 373, eabf3002. 10.1126/science.abf3002.

24. Naik, S., Larsen, S.B., Gomez, N.C., Alaverdyan, K., Sendoel, A., Yuan, S., Polak, L., Kulukian, A., Chai, S., and Fuchs, E. (2017). Inflammatory memory sensitizes skin epithelial stem cells to tissue damage. Nature 550, 475–480. 10.1038/nature24271.

25. Larsen, S.B., Cowley, C.J., Sajjath, S.M., Barrows, D., Yang, Y., Carroll, T.S., and Fuchs, E. (2021). Establishment, maintenance, and recall of inflammatory memory. Cell Stem Cell 28, 1758–1774.e8. 10.1016/j.stem.2021.07.001.

26. Gonzales, K.A.U., Polak, L., Matos, I., Tierney, M.T., Gola, A., Wong, E., Infarinato, N.R., Nikolova, M., Luo, S., Liu, S., et al. (2021). Stem cells expand potency and alter tissue fitness by accumulating diverse epigenetic memories. Science 374, eabh2444. 10.1126/science.abh2444.

27. Ordovas-Montanes, J., Dwyer, D.F., Nyquist, S.K., Buchheit, K.M., Vukovic, M., Deb, C., Wadsworth, M.H., Hughes, T.K., Kazer, S.W., Yoshimoto, E., et al. (2018). Allergic inflammatory memory in human respiratory epithelial progenitor cells. Nature 560, 649–654. 10.1038/s41586-018-0449-8.

28. Fenton, C.G., Taman, H., Florholmen, J., Sørbye, S.W., and Paulssen, R.H. (2021). Transcriptional Signatures That Define Ulcerative Colitis in Remission. Inflamm Bowel Dis 27, 94–105. 10.1093/ibd/izaa075.

29. Planell, N., Lozano, J.J., Mora-Buch, R., Masamunt, M.C., Jimeno, M., Ordás, I., Esteller, M., Ricart, E., Piqué, J.M., Panés, J., et al. (2013). Transcriptional analysis of the intestinal mucosa of patients with ulcerative colitis in remission reveals lasting epithelial cell alterations. Gut 62, 967–976. 10.1136/gutjnl-2012-303333.

30. Howell, K.J., Kraiczy, J., Nayak, K.M., Gasparetto, M., Ross, A., Lee, C., Mak, T.N., Koo, B.-K., Kumar, N., Lawley, T., et al. (2018). DNA Methylation and Transcription Patterns in Intestinal Epithelial Cells From Pediatric Patients With Inflammatory Bowel Diseases Differentiate Disease Subtypes and Associate With Outcome. Gastroenterology 154, 585–598. 10.1053/j.gastro.2017.10.007.

31. Singh, V., Johnson, K., Yin, J., Lee, S., Lin, R., Yu, H., In, J., Foulke-Abel, J., Zachos, N.C., Donowitz, M., et al. (2022). Chronic Inflammation in Ulcerative Colitis Causes Long-Term Changes in Goblet Cell Function. Cell Mol Gastroenterol Hepatol 13, 219–232. 10.1016/j.jcmgh.2021.08.010.

32. JR, K., N, D., MA, C., TA, K., K, M., JM, W., S, U., MH, D., R, B., LM, B., et al. (2021). Colonoids From Patients With Pediatric Inflammatory Bowel Disease Exhibit Decreased Growth Associated With Inflammation Severity and Durable Upregulation of Antigen Presentation Genes. Inflammatory bowel diseases 27, 256–267. 10.1093/IBD/IZAA145.

33. Zhao, D., Ravikumar, V., Leach, T.J., Kraushaar, D., Lauder, E., Li, L., Sun, Y., Oravecz-Wilson, K., Keller, E.T., Chen, F., et al. (2024). Inflammation-induced epigenetic imprinting regulates intestinal stem cells. Cell Stem Cell, S1934–5909(24)00292-3. 10.1016/j.stem.2024.08.006.

34. Morral, C., Ghinnagow, R., Karakasheva, T., Zhou, Y., Thadi, A., Li, N., Yoshor, B., Soto, G.E., Chen, C.-H., Aleynick, D., et al. (2023). Isolation of Epithelial and Stromal Cells from Colon Tissues in Homeostasis and Under Inflammatory Conditions. Bio Protoc 13, e4825. 10.21769/BioProtoc.4825.

35. Asaf, S., Maqsood, F., Jalil, J., Sarfraz, Z., Sarfraz, A., Mustafa, S., and Ojeda, I.C. (2023). Lipocalin 2-not only a biomarker: a study of current literature and systematic findings of ongoing clinical trials. Immunol Res 71, 287–313. 10.1007/s12026-022-09352-2.

36. Toyonaga, T., Matsuura, M., Mori, K., Honzawa, Y., Minami, N., Yamada, S., Kobayashi, T., Hibi, T., and Nakase, H. (2016). Lipocalin 2 prevents intestinal inflammation by enhancing phagocytic bacterial clearance in macrophages. Sci Rep 6, 35014. 10.1038/srep35014.

37. Moschen, A.R., Gerner, R.R., Wang, J., Klepsch, V., Adolph, T.E., Reider, S.J., Hackl, H., Pfister, A., Schilling, J., Moser, P.L., et al. (2016). Lipocalin 2 Protects from Inflammation and Tumorigenesis Associated with Gut Microbiota Alterations. Cell Host Microbe 19, 455–469. 10.1016/j.chom.2016.03.007.

38. Lawrance, I.C. (2001). Ulcerative colitis and Crohn’s disease: distinctive gene expression profiles and novel susceptibility candidate genes. Human Molecular Genetics 10, 445–456. 10.1093/hmg/10.5.445.

39. Renner, M., Bergmann, G., Krebs, I., End, C., Lyer, S., Hilberg, F., Helmke, B., Gassler, N., Autschbach, F., Bikker, F., et al. (2007). DMBT1 Confers Mucosal Protection In Vivo and a Deletion Variant Is Associated With Crohn’s Disease. Gastroenterology 133, 1499–1509. 10.1053/j.gastro.2007.08.007.

40. James, J.P., Nielsen, B.S., Christensen, I.J., Langholz, E., Malham, M., Poulsen, T.S., Holmstrøm, K., Riis, L.B., and Høgdall, E. (2023). Mucosal expression of PI3, ANXA1, and VDR discriminates Crohn’s disease from ulcerative colitis. Sci Rep 13, 18421. 10.1038/s41598-023-45569-3.

41. Farr, L., Ghosh, S., Jiang, N., Watanabe, K., Parlak, M., Bucala, R., and Moonah, S. (2020). CD74 Signaling Links Inflammation to Intestinal Epithelial Cell Regeneration and Promotes Mucosal Healing. Cell Mol Gastroenterol Hepatol 10, 101–112. 10.1016/j.jcmgh.2020.01.009.

42. Stokkers, P., Reitsma, P., Tytgat, G., and Deventer, S.J.H. van (1999). HLA-DR and -DQ phenotypes in inflammatory bowel disease: a meta-analysis. Gut 45, 395. 10.1136/gut.45.3.395.

43. HLA-DP on Epithelial Cells Enables Tissue Damage by NKp44+ Natural Killer Cells in Ulcerative Colitis - Gastroenterology https://www.gastrojournal.org/article/S0016-5085(23)04772-8/fulltext.

44. Grasberger, H., Magis, A.T., Sheng, E., Conomos, M.P., Zhang, M., Garzotto, L.S., Hou, G., Bishu, S., Nagao-Kitamoto, H., El-Zaatari, M., et al. (2021). DUOX2 variants associate with preclinical disturbances in microbiota-immune homeostasis and increased inflammatory bowel disease risk. J Clin Invest 131, e141676, 141676. 10.1172/JCI141676.

45. Stettner, N., Rosen, C., Bernshtein, B., Gur-Cohen, S., Frug, J., Silberman, A., Sarver, A., Carmel-Neiderman, N.N., Eilam, R., Biton, I., et al. (2018). Induction of Nitric-Oxide Metabolism in Enterocytes Alleviates Colitis and Inflammation-Associated Colon Cancer. Cell Reports 23, 1962. 10.1016/j.celrep.2018.04.053.

46. Schewe, M., Franken, P.F., Sacchetti, A., Schmitt, M., Joosten, R., Böttcher, R., van Royen, M.E., Jeammet, L., Payré, C., Scott, P.M., et al. (2016). Secreted Phospholipases A2 Are Intestinal Stem Cell Niche Factors with Distinct Roles in Homeostasis, Inflammation, and Cancer. Cell Stem Cell 19, 38–51. 10.1016/j.stem.2016.05.023.

47. Kaser, A., Ludwiczek, O., Holzmann, S., Moschen, A.R., Weiss, G., Enrich, B., Graziadei, I., Dunzendorfer, S., Wiedermann, C.J., Mürzl, E., et al. (2004). Increased expression of CCL20 in human inflammatory bowel disease. J Clin Immunol 24, 74–85. 10.1023/B:JOCI.0000018066.46279.6b.

48. Marafini, I., Monteleone, I., Dinallo, V., Di Fusco, D., De Simone, V., Laudisi, F., Fantini, M.C., Di Sabatino, A., Pallone, F., and Monteleone, G. (2016). CCL20 Is Negatively Regulated by TGF-β1 in Intestinal Epithelial Cells and Reduced in Crohn’s Disease Patients With a Successful Response to Mongersen, a Smad7 Antisense Oligonucleotide. ECCOJC, jjw191. 10.1093/ecco-jcc/jjw191.

49. Baumdick, M.E., Niehrs, A., Degenhardt, F., Schwerk, M., Hinrichs, O., Jordan-Paiz, A., Padoan, B., Wegner, L.H.M., Schloer, S., Zecher, B.F., et al. (2023). HLA-DP on Epithelial Cells Enables Tissue Damage by NKp44+ Natural Killer Cells in Ulcerative Colitis. Gastroenterology 165, 946–962.e13. 10.1053/j.gastro.2023.06.034.

50. Schumacher, M.A. (2023). The emerging roles of deep crypt secretory cells in colonic physiology. American Journal of Physiology-Gastrointestinal and Liver Physiology 325, G493–G500. 10.1152/ajpgi.00093.2023.

51. Sasaki, N., Sachs, N., Wiebrands, K., Ellenbroek, S.I., Fumagalli, A., Lyubimova, A., Begthel, H., van den Born, M., van Es, J.H., Karthaus, W.R., et al. (2016). Reg4+ deep crypt secretory cells function as epithelial niche for Lgr5+ stem cells in colon. Proc Natl Acad Sci U S A 113, E5399–407. 10.1073/pnas.1607327113.

52. Hyams, J.S., Ferry, G.D., Mandel, F.S., Gryboski, J.D., Kibort, P.M., Kirschner, B.S., Griffiths, A.M., Katz, A.J., Grand, R.J., and Boyle, J.T. (1991). Development and validation of a pediatric Crohn’s disease activity index. J Pediatr Gastroenterol Nutr 12, 439–447.

53. Hyams, J., Crandall, W., Kugathasan, S., Griffiths, A., Olson, A., Johanns, J., Liu, G., Travers, S., Heuschkel, R., Markowitz, J., et al. (2007). Induction and Maintenance Infliximab Therapy for the Treatment of Moderate-to-Severe Crohn’s Disease in Children. Gastroenterology 132, 863–873. 10.1053/j.gastro.2006.12.003.

54. Ayyaz, A., Kumar, S., Sangiorgi, B., Ghoshal, B., Gosio, J., Ouladan, S., Fink, M., Barutcu, S., Trcka, D., Shen, J., et al. (2019). Single-cell transcriptomes of the regenerating intestine reveal a revival stem cell. Nature 2019 569:7754 569, 121–125. 10.1038/s41586-019-1154-y.

55. Sprangers, J., Zaalberg, I.C., and Maurice, M.M. (2021). Organoid-based modeling of intestinal development, regeneration, and repair. Cell Death Differ 28, 95–107. 10.1038/s41418-020-00665-z.

56. Qiao, X.T., Ziel, J.W., McKimpson, W., Madison, B.B., Todisco, A., Merchant, J.L., Samuelson, L.C., and Gumucio, D.L. (2007). Prospective Identification of a Multilineage Progenitor in Murine Stomach Epithelium. Gastroenterology 133, 1989–1998.e3. 10.1053/j.gastro.2007.09.031.

57. Burclaff, J., Bliton, R.J., Breau, K.A., Ok, M.T., Gomez-Martinez, I., Ranek, J.S., Bhatt, A.P., Purvis, J.E., Woosley, J.T., and Magness, S.T. (2022). A Proximal-to-Distal Survey of Healthy Adult Human Small Intestine and Colon Epithelium by Single-Cell Transcriptomics. CMGH 13, 1554–1589. 10.1016/j.jcmgh.2022.02.007.

58. Cheong, J.-G., Ravishankar, A., Sharma, S., Parkhurst, C.N., Grassmann, S.A., Wingert, C.K., Laurent, P., Ma, S., Paddock, L., Miranda, I.C., et al. (2023). Epigenetic memory of coronavirus infection in innate immune cells and their progenitors. Cell 186, 3882–3902.e24. 10.1016/j.cell.2023.07.019.

59. Funk, M.C., Gleixner, J.G., Heigwer, F., Vonficht, D., Valentini, E., Aydin, Z., Tonin, E., Prete, S.D., Mahara, S., Throm, Y., et al. (2023). Aged intestinal stem cells propagate cell-intrinsic sources of inflammaging in mice. Developmental Cell 58, 2914–2929.e7. 10.1016/j.devcel.2023.11.013.

60. Adegbola, S.O., Sahnan, K., Warusavitarne, J., Hart, A., and Tozer, P. (2018). Anti-TNF Therapy in Crohn’s Disease. International Journal of Molecular Sciences 19, 2244. 10.3390/ijms19082244.

61. Arnauts, K., Verstockt, B., Ramalho, A.S., Vermeire, S., Verfaillie, C., and Ferrante, M. (2020). Ex Vivo Mimicking of Inflammation in Organoids Derived From Patients With Ulcerative Colitis. Gastroenterology 159, 1564–1567. 10.1053/j.gastro.2020.05.064.

62. d’Aldebert, E., Quaranta, M., Sébert, M., Bonnet, D., Kirzin, S., Portier, G., Duffas, J.P., Chabot, S., Lluel, P., Allart, S., et al. (2020). Characterization of Human Colon Organoids From Inflammatory Bowel Disease Patients. Frontiers in Cell and Developmental Biology 8, 363. 10.3389/FCELL.2020.00363/BIBTEX.

63. Podolsky, D.K. (2002). Inflammatory Bowel Disease. New England Journal of Medicine 347, 417–429. 10.1056/NEJMra020831.

64. Jamwal, D.R., Laubitz, D., Harrison, C.A., Figliuolo da Paz, V., Cox, C.M., Wong, R., Midura-Kiela, M., Gurney, M.A., Besselsen, D.G., Setty, P., et al. (2020). Intestinal Epithelial Expression of MHCII Determines Severity of Chemical, T-Cell-Induced, and Infectious Colitis in Mice. Gastroenterology 159, 1342–1356.e6. 10.1053/j.gastro.2020.06.049.

65. Kyodo, R., Takeuchi, I., Narumi, S., Shimizu, H., Hata, K., Yoshioka, T., Tanase-Nakao, K., Shimizu, T., and Arai, K. (2022). Novel biallelic mutations in the DUOX2 gene underlying very early-onset inflammatory bowel disease: A case report. Clin Immunol 238, 109015. 10.1016/j.clim.2022.109015.

66. Hayes, P., Dhillon, S., O’Neill, K., Thoeni, C., Hui, K.Y., Elkadri, A., Guo, C.H., Kovacic, L., Aviello, G., Alvarez, L.A., et al. (2015). Defects in NADPH Oxidase Genes NOX1 and DUOX2 in Very Early Onset Inflammatory Bowel Disease. Cell Mol Gastroenterol Hepatol 1, 489–502. 10.1016/j.jcmgh.2015.06.005.

67. Hazime, H., Ducasa, G.M., Santander, A.M., Brito, N., González, E.E., Ban, Y., Kaunitz, J., Akiba, Y., Fernández, I., Burgueño, J.F., et al. (2023). Intestinal Epithelial Inactivity of Dual Oxidase 2 Results in Microbiome-Mediated Metabolic Syndrome. Cell Mol Gastroenterol Hepatol 16, 557–572. 10.1016/j.jcmgh.2023.06.009.

68. 68. Liu, Z., Cominelli, F., Di Martino, L., Liu, R., Devireddy, N., Devireddy, L.R., and Wald, D.N. (2019). Lipocalin 24p3 Induction in Colitis Adversely Affects Inflammation and Contributes to Mortality. Front Immunol 10, 812. 10.3389/fimmu.2019.00812.

69. La Manno, G., Soldatov, R., Zeisel, A., Braun, E., Hochgerner, H., Petukhov, V., Lidschreiber, K., Kastriti, M.E., Lönnerberg, P., Furlan, A., et al. (2018). RNA velocity of single cells. Nature 560, 494–498. 10.1038/s41586-018-0414-6.

70. Li, J., Simmons, A.J., Hawkins, C.V., Chiron, S., Ramirez-Solano, M.A., Tasneem, N., Kaur, H., Xu, Y., Revetta, F., Vega, P.N., et al. (2024). Identification and multimodal characterization of a specialized epithelial cell type associated with Crohn’s disease. Nat Commun 15, 7204. 10.1038/s41467-024-51580-7.

71. Schumacher, M.A., Liu, C.Y., Katada, K., Thai, M.H., Hsieh, J.J., Hansten, B.J., Waddell, A., Rosen, M.J., and Frey, M.R. (2023). Deep Crypt Secretory Cell Differentiation in the Colonic Epithelium Is Regulated by Sprouty2 and Interleukin 13. Cellular and Molecular Gastroenterology and Hepatology 15, 971–984. 10.1016/j.jcmgh.2022.11.004.

72. Liu, C.Y., Girish, N., Gomez, M.L., Kalski, M., Bernard, J.K., Simons, B.D., and Polk, D.B. (2023). Wound-healing plasticity enables clonal expansion of founder progenitor cells in colitis. Dev Cell 58, 2309–2325.e7. 10.1016/j.devcel.2023.08.011.

73. Verhagen, M.P., Joosten, R., Schmitt, M., Välimäki, N., Sacchetti, A., Rajamäki, K., Choi, J., Procopio, P., Silva, S., van der Steen, B., et al. (2024). Non-stem cell lineages as an alternative origin of intestinal tumorigenesis in the context of inflammation. Nat Genet 56, 1456–1467. 10.1038/s41588-024-01801-y.

74. Zhao, D., Ravikumar, V., Leach, T.J., Kraushaar, D., Lauder, E., Li, L., Sun, Y., Oravecz-Wilson, K., Keller, E.T., Chen, F., et al. (2024). Inflammation-induced epigenetic imprinting regulates intestinal stem cells. Cell Stem Cell, S1934–5909(24)00292-3. 10.1016/j.stem.2024.08.006.

75. Matsuzawa-Ishimoto, Y., Hine, A., Shono, Y., Rudensky, E., Lazrak, A., Yeung, F., Neil, J.A., Yao, X., Chen, Y.-H., Heaney, T., et al. (2020). An intestinal organoid-based platform that recreates susceptibility to T-cell-mediated tissue injury. Blood 135, 2388–2401. 10.1182/blood.2019004116.

76. Takashima, S., Martin, M.L., Jansen, S.A., Fu, Y., Bos, J., Chandra, D., O’Connor, M.H., Mertelsmann, A.M., Vinci, P., Kuttiyara, J., et al. (2019). T cell-derived interferon-γ programs stem cell death in immune-mediated intestinal damage. Sci Immunol 4, eaay8556. 10.1126/sciimmunol.aay8556.

77. Bolstad, B. (2024). bmbolstad/preprocessCore.

78. Becht, E., McInnes, L., Healy, J., Dutertre, C.-A., Kwok, I.W.H., Ng, L.G., Ginhoux, F., and Newell, E.W. (2018). Dimensionality reduction for visualizing single-cell data using UMAP. Nat Biotechnol. 10.1038/nbt.4314.

79. Qiu, X., Mao, Q., Tang, Y., Wang, L., Chawla, R., Pliner, H.A., and Trapnell, C. (2017). Reversed graph embedding resolves complex single-cell trajectories. Nat Methods 14, 979–982. 10.1038/nmeth.4402.

80. Gao, M., Qiao, C., and Huang, Y. (2022). UniTVelo: temporally unified RNA velocity reinforces single-cell trajectory inference. Nat Commun 13, 6586. 10.1038/s41467-022-34188-7.

81. Weiler, P., Lange, M., Klein, M., Pe’er, D., and Theis, F. (2024). CellRank 2: unified fate mapping in multiview single-cell data. Nat Methods 21, 1196–1205. 10.1038/s41592-024-02303-9.

82. Karakasheva, T. (2023). Working with patient-derived enteroids and colonoids. 10.17504/protocols.io.ewov1q7e2gr2/v1.

83. Fujii, M., Matano, M., Toshimitsu, K., Takano, A., Mikami, Y., Nishikori, S., Sugimoto, S., and Sato, T. (2018). Human Intestinal Organoids Maintain Self-Renewal Capacity and Cellular Diversity in Niche-Inspired Culture Condition. Cell stem cell 23, 787–793.e6. 10.1016/J.STEM.2018.11.016.

84. VanDussen, K.L., Sonnek, N.M., and Stappenbeck, T.S. (2019). L-WRN conditioned medium for gastrointestinal epithelial stem cell culture shows replicable batch-to-batch activity levels across multiple research teams. Stem Cell Research 37, 101430. 10.1016/j.scr.2019.101430.

